# Mapping of the podocin proximity-dependent proteome reveals novel components of the kidney podocyte foot process

**DOI:** 10.1101/2022.11.03.515102

**Authors:** Gary F. Gerlach, Zachary H. Imseis, Shamus L. Cooper, Anabella N. Santos, Lori L. O’Brien

**Affiliations:** Department of Cell Biology and Physiology, University of North Carolina at Chapel Hill, Chapel Hill, NC 27599

## Abstract

The unique architecture of glomerular podocytes is integral to kidney filtration. Interdigitating foot processes extend from the podocyte cell body, wrap around fenestrated capillaries, and form specialized junctional complexes termed slit diaphragms to create a molecular sieve. However, the full complement of proteins which maintain foot process integrity, and how this localized proteome changes with disease, remains to be elucidated. Proximity-dependent <u>bio</u>tin <u>id</u>entification (BioID) enables the identification of spatially localized proteomes. To this end, we developed a novel *in vivo* BioID knock-in mouse model. We utilized the slit diaphragm protein podocin (*Nphs2*) to create a podocin-BioID fusion. Podocin-BioID localizes to the slit diaphragm and biotin injection leads to podocyte-specific protein biotinylation. We isolated the biotinylated proteins and performed mass spectrometry to identify proximal interactors. Gene ontology analysis of 54 proteins specifically enriched in our podocin-BioID sample revealed ‘cell junctions’, ‘actin binding’, and ‘cytoskeleton organization’ as top terms. Known foot process components were identified and we further uncovered two novel proteins: the tricellular junctional protein Ildr2 and the CDC42 and N-WASP interactor Fnbp1l. We confirmed Ildr2 and Fnbp1l are expressed by podocytes and partially colocalize with podocin. Finally, we investigated how this proteome changes with age and uncovered a significant increase in Ildr2. This was confirmed by immunofluorescence on human kidney samples and suggests altered junctional composition may preserve podocyte integrity. Together, these assays have led to new insights into podocyte biology and supports the efficacy of utilizing BioID *in vivo* to interrogate spatially localized proteomes in health, aging, and disease.

## Introduction

Kidneys perform vital functions as they filter waste and toxins from the blood and regulate body fluid homeostasis. The simplest functional unit of the kidney is the nephron, composed of a blood filter termed the glomerulus connected to a segmented tubule. The glomerulus operates as a ‘molecular sieve’, filtering blood and inhibiting passage of large macromolecules and red blood cells into the nephron tubule. The glomerulus relies on specialized epithelial cells called podocytes, named for their unique cellular morphology with long extruding projections, termed foot processes. Podocyte foot processes wrap around a tuft of fenestrated endothelial capillaries leaving small gaps, or slits, between them. Podocytes undergo morphological changes to their junctional architecture during development to form a specialized barrier between foot processes termed a slit diaphragm.^1–3^ The slit diaphragm executes multiple functions including macromolecular filtering in collaboration with the underlying glomerular basement membrane, connection to the actin cytoskeleton to maintain foot process architecture, and signaling that regulates podocyte integrity. The slit diaphragm is distinct from other junctional complexes as it integrates unique structural components as well as components of adherens and tight junctions ^4–8^. Nearly 50 years ago, a zipper-like model for the slit diaphragm was proposed, wherein proteins from neighboring foot processes partially cross the intervening intercellular space and overlap, forming the dense protein-rich slit diaphragm structure eloquently visualized by electron microscopy (EM).^9,10^ More recent block-face scanning electron microscopy has revealed a ‘ridge-like prominence’ architecture to podocyte foot processes, formed on the basal surface of the primary foot process.^2,11^ These investigations underscore the continued advancement in our understanding of podocyte structure and function.

When podocytes undergo stress or injury above a threshold, they initiate a response that leads to foot process effacement, loss of slit diaphragms, and proteinuria. Loss of podocyte integrity, observed as effacement, is associated with proteinuric kidney disease^12–14^. This pathology has been described in both acquired and hereditary forms of glomerular disorders or podocytopathies.^15^ Podocytopathies are a class of kidney diseases in which direct or indirect podocyte injury drives proteinuria or nephrotic syndrome and can ultimately lead to end-stage renal disease (ESRD). Genetic studies have previously identified mutations in numerous podocyte foot process components such as Membrane associated guanylate kinase WW and PDZ Domain Containing 2 (MAGI2)^16^, CD2-associated protein (CD2AP)^17,18^, nephrin (NPHS1)^6,19^, and podocin (NPHS2)^7,20^ as causal for nephrotic disease. Additionally, diseases such as diabetes and autoimmune disorders can lead to podocyte injury^21^. While the downstream result is effacement and loss of slit diaphragms, we know little about the temporal changes occurring specifically within the foot process and locally at the slit diaphragm.

The identification of slit diaphragm protein complexes with immunoprecipitation followed by mass spectrometry (MS) has uncovered important localized interactions^22,23^. However, the efficiency of these immunoprecipitations is often hindered by harsh conditions required to extract membrane proteins, which can eliminate weaker binding interactions, or by antibodies that may disrupt interactions. Additionally, transient interactions may be missed in such experiments. Podocin is known to localize to the slit diaphragm and interacts with Nphs1 (nephrin), Neph1 (Kirrel), and Cd2ap.^24–27^ Podocin is a member of the stomatin family, containing a central hinge region that integrates into the membrane of the foot process with cytoplasmic N and C termini.^28^ Previous studies demonstrated podocin’s role in the development of the multiprotein-lipid super complex of the slit diaphragm.^27^ Podocin’s ability to oligomerize and act as a protein scaffold at slit diaphragms ideally positions it for use as a bait protein in proteomic studies.

The discovery and engineering of a promiscuous prokaryotic biotin ligase by Roux and colleagues and concomitant blossoming of –omics technologies in the last decade have laid the groundwork to uncover spatially localized proteomes.^29,30^ Proximity-dependent biotin identification, or BioID, utilizes a mutated prokaryotic biotin ligase fused to a bait protein of interest to covalently attach biotin to proteins within the vicinity of a bait protein. The radius of biotinylation can range from ∼10nm to 25nm, dependent on the size of the linker between the bait and the biotin ligase, allowing for the biotinylation of both direct and indirect interactors.^29,30^ The BioID system requires exogenous addition of biotin for the ligase to covalently biotinylate a proximally located protein, giving the BioID system spatiotemporal control of protein tagging. Additionally, the biotin labels are stable and can withstand harsh isolation conditions, allowing the capture of transient interactors and membrane proteins, respectively. The BioID system has provided plentiful *in vitro* reports from cell culture models that highlight the power of the system and its ability to discover novel components of even well-documented cellular machinery such as the centrosome and cilium.^31,32^ There have been limited *in vivo* reports of BioID, but it has been used successfully in vertebrates such as zebrafish and mice to identify endogenous interactomes.^32–34^ To the best of our knowledge, this approach has not been employed in the mammalian kidney.

Here, we utilized gene editing to introduce a smaller, more efficient promiscuous biotin ligase (BioID2) into the endogenous murine *Nphs2* locus to create a fusion protein, hereafter referred to as podocin-BioID (*Nphs2*^*BioID2*^). Our podocin-BioID model offers the capacity to uncover proteins that localize to the region of the podocyte foot process within the vicinity of podocin in an *in vivo* mammalian system. We were able to identify novel podocyte foot process proteins and furthermore how this proteome changes with age, highlighting the utility of our model for uncovering new interactors as well as disease-associated changes.

## Results

### Generation of the *Nphs2*^*BioID2*^ knock-in mouse model

To generate our mouse model, we utilized CRISPR/Cas9 gene editing in combination with homology-directed repair to knock-in the HA-tagged, mutated *A. aeolicus* biotin ligase (BioID2) in frame at the *Nphs2* locus^30^. A single guide RNA (sgRNA) was used to target the stop codon within the eighth exon of *Nphs2* to create the fusion. A 13x Glycine/Serine(G/S) flexible linker region was included between *Nphs2* and the ligase to provide up to a 25nm reach (Figure 1A)^30^. C57BL/6J zygotes were injected and after screening the resulting animals for a single knock-in with the correct sequence, several founders were identified. A single male founder was utilized for subsequent breeding and expansion of the *Nphs2*^*BioID2*^ line. Genotyping E18.5-P0 pups from incrosses of *Nphs2*^*BioID2/+*^ animals identified wildtype, heterozygous, and homozygous offspring at approximately anticipated Mendelian ratios of 29%, 52%, and 19%, respectively (n=73, Figure 1B,C). However, we were typically unable to recover homozygous *Nphs2*^*BioID2/BioID2*^ pups after 1-week of birth. Overall, the gross morphology of homozygous pups and kidneys appeared normal at P0 and we were unable to determine the specific cause of their death. Due to the early homozygous death, all subsequent experiments for proteomic profiling were performed on heterozygous *Nphs2*^*BioID2/+*^ animals. To test the function of the podocin-BioID fusion and determine if the ligase was able to biotinylate podocyte proteins, we administered 5 mg/kg biotin for 7 consecutive days to 8-10 week old *Nphs2*^*BioID2/+*^ and wild type control mice, a dosage utilized in previous protocols for *in vivo* BioID experiments.^34^ Following 1 week of subcutaneous biotin injections, kidneys harvested from *Nphs2*^*BioID2/+*^ mice displayed a pronounced streptavidin signal within glomeruli and specifically podocytes marked by Wt1-positive nuclei (Figure 1D). We occasionally detected streptavidin-positive signal within the tubules which either represents background staining or uptake of free biotin, as controls also displayed this non-glomerular signal pattern. Finally, probing for the HA tag contained within the podocin-BioID fusion shows that the HA signal overlaps with the streptavidin signal, confirming the fusion protein is being specifically translated in podocytes and that the biotin ligase is functional (Figure 1D).

**Figure 1.**
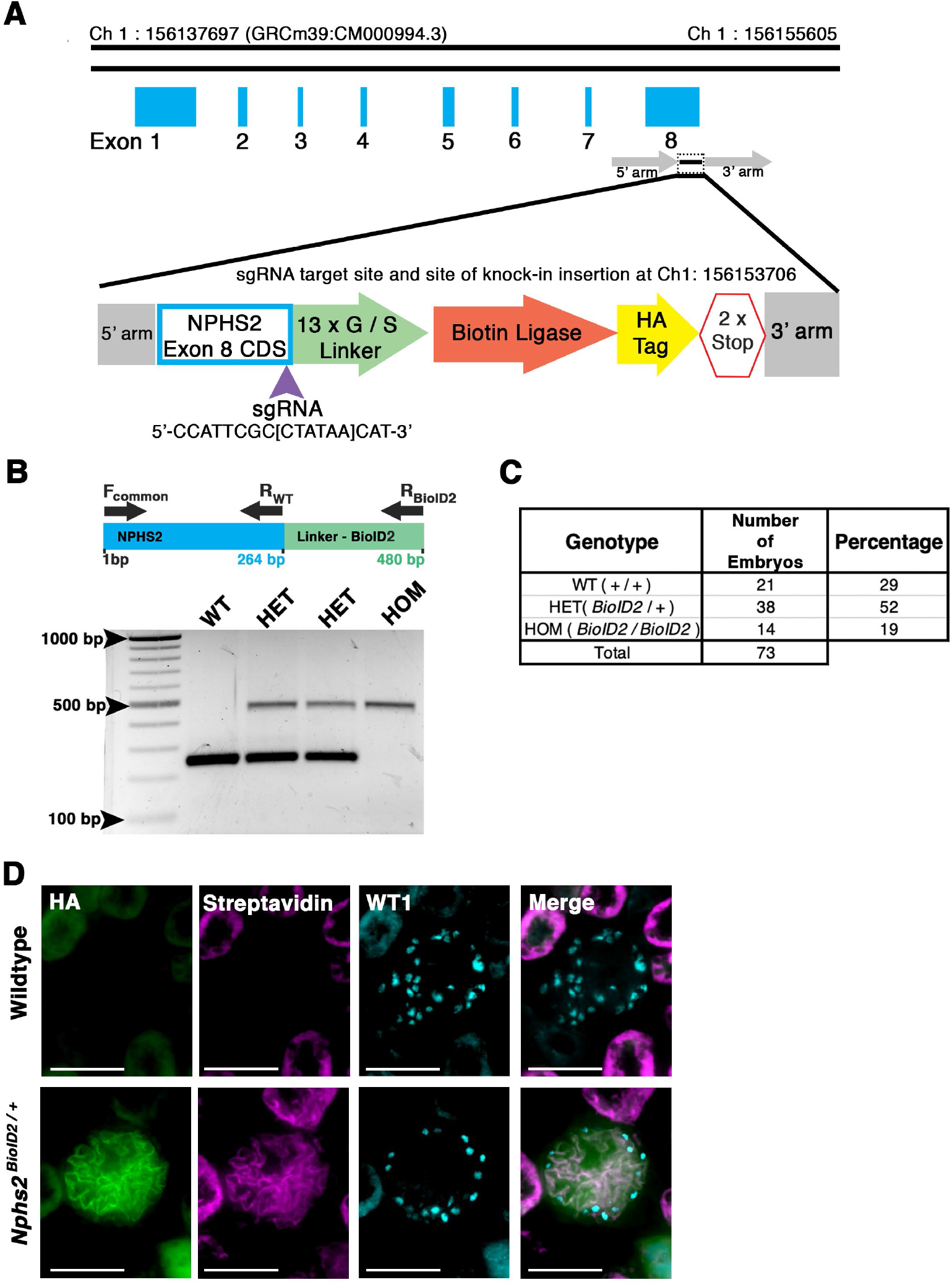
Generation of a knock-in *Nphs2*^*BioID2*^ mouse line via CRISPR/Cas9 genome editing of the *Nphs2* locus. **(A)** Schematic of the CRISPR/Cas9 genome editing strategy utilized to generate the *Nphs2*^*BioID2*^ mouse line. A small guide RNA (sgRNA) targeting the stop codon in exon 8 of *Nphs2* (purple arrowhead) was combined with a donor vector containing the knock-in cassette (zoom view) to induce homologous recombination and integrate the BioID2 moiety containing a 13x Glycine/Serine (G/S) linker, biotin ligase, and HA tag into the *Nphs2* locus. **(B)** Genotyping strategy (top panel). Wildtype (single band at 264 base pairs (bp)), heterozygous (two bands; one at wildtype size of 264 bp and a second that amplifies the BioID2 linker region giving a band at 480 bp), or homozygous (single band at the 480 bp). **(C)** Genotyping of 73 embryos at E18.5-P0 verify a relative Mendelian ratio of genotypes being recovered (25:50:25). **(D)** Immunofluorescence analysis of 8–10-week-old adult mice injected with biotin show an enrichment of streptavidin within the glomerulus of *Nphs2*^*BioID2*^ mice and absence of streptavidin signal in control littermate mice. The HA signal from the BioID2 moiety closely overlaps with streptavidin (merged image). Wt1 (cyan) is utilized as a podocyte marker to delineate the glomerular boundaries. Scale bar: 50 μm

### Podocin-BioID kidneys display normal nephron morphology and the fusion localizes to the slit diaphragm

We further wanted to confirm that the kidneys of animals expressing the podocin-BioID fusion did not display any significant phenotypic differences, most specifically to the nephron. If the fusion protein was not localizing or functioning properly, the animals may display phenotypes associated with *Nphs2* knockout animals such as enlarged glomeruli, vacuolated podocytes, and mesangial expansion.^35^ Additionally, dilated tubules may indicate abnormal nephron function. Immunostaining kidney sections of wild type, heterozygous, and homozygous animals at E18.5 revealed no qualitative differences in glomerular size or proximal tubule dilation (Figure 2A). This further supports that mislocalization or abnormal function of podocin-BioID in podocytes is not the likely cause of death in the homozygous animals. Immunofluorescence staining showed a significant overlap between the HA signal and signal from a podocin-specific antibody, supporting that the fusion protein is localizing properly in heterozygous animals and that it is being expressed at similar levels in the homozygous animals (Figure 2A). Importantly, we did not observe the HA antibody signal anywhere besides the glomerulus.

**Figure 2.**
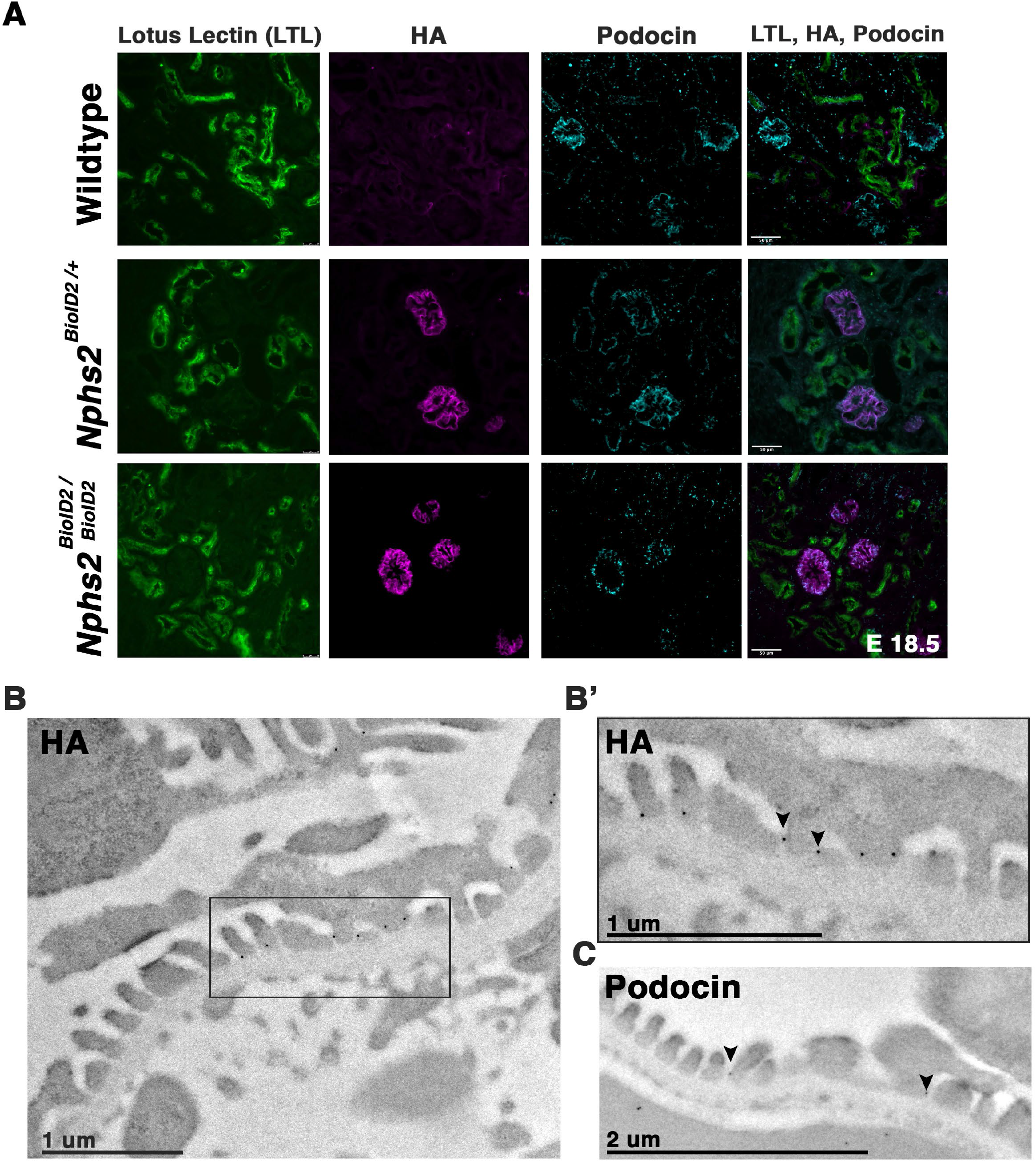
*Nphs2*^*BioID2*^ animals present normal kidney morphology with podocin-BioID localized to podocytes and specifically the slit diaphragm. **(A)** E18.5 littermates of wildtype, heterozygous (*Nphs2*^*BioID2/+*^), and homozygous (*Nphs2*^*BioID2/BioID2*^) animals were analyzed for gross nephron morphology and localization of podocin-BioID. Proximal tubule: (LTL, green); BioID: (HA, magenta); podocytes/glomerulus: (podocin, cyan). Scale bar: 50 μm **(B)** Transmission electron microscopy (TEM) and immunogold labeling of the HA (BioID) of 4-week-old *Nphs2*^*BioID2/+*^ kidneys. Boxed area is shown in B’. **(B’)** Magnified view of boxed region from panel (B). HA signal localizes as dark spherical dots from immunogold labeling (arrowheads). Scale bar: 1 μm **(C)** Wildtype littermates immunogold labeled for podocin (arrowheads). Scale bar: 2 μm.

Additionally, we wanted to confirm that the podocin-BioID protein was localizing to the slit diaphragm where podocin is known to interact with other slit diaphragm proteins and play a functional role.^24–27,35^ We utilized immunogold labeling in combination with transmission electron microscopy (TEM) to examine the subcellular localization of the podocin-BioID protein (Figure 2B,B’). *Nphs2*^*BioID2+*^ kidney sections were stained with an anti-HA antibody followed by a colloidal gold-AffiniPure secondary. Punctate gold signals were observed near electron dense regions between foot processes, the location of the slit diaphragm (Figure 2B,B’ arrowheads). We observed minimal gold signal outside of this region. To confirm that this is the site of normal podocin localization in our control animals, wildtype littermates were probed with a podocin antibody and also showed localization of punctate gold signal near the electron dense region of the slit diaphragm (Figure 2C, arrowheads).

### An enrichment of biotinylated proteins is detected in lysates from *Nphs2*^*BioID2/+*^ kidneys

To enrich our podocin-BioID protein lysates for glomeruli, we surgically isolated the cortex from each kidney of biotin injected *Nphs2*^*BioID2/+*^ animals and wildtype controls at 8-10 weeks. Isolated cortex was homogenized and lysed to obtain protein lysates from each animal. Protein lysates were applied to magnetic streptavidin coated beads to isolate the biotinylated proteins and an aliquot was removed and tested to validate the efficacy of the biotin ligase. By Western blot analysis, we identified an increase in the number of biotinylated proteins in our *Nphs2*^*BioID2/+*^ sample versus wildtype along the full spectrum of molecular weights (Figure 3A). In contrast, few streptavidin labeled, biotinylated, proteins were visible in controls. The few biotinylated proteins observed in wildtype littermates likely represent the endogenous metabolic CoA carboxylases^36^. Due to podocin’s ability to oligomerize and the heterozygous nature of the mice, we would expect that endogenous, non-tagged podocin would be biotinylated as well as podocin-BioID itself. When we probed the Western blot for podocin, we observed two bands within the *Nphs2*^*BioID2/+*^ sample: one band at approximately 50 kDa, the predicted endogenous podocin molecular weight without the BioID2 tag (Figure 3C, denoted with an asterisk), and a second larger protein, podocin-BioID (Figure 3C, denoted with arrowhead). No bands for podocin or podocin-BioID were detected in controls, as expected. Additionally, the HA signal on the BioID2 protein was only detected in our *Nphs2*^*BioID2/+*^ protein lysates confirming the purity and specificity of our results (Figure 3B). We went on to test if biotin could cross the placental barrier to be delivered to embryonic pups *via* injection of the pregnant dam. Pregnant dams were injected each day for one week, from E11.5-E18.5, subcutaneously with 5 mg/kg biotin. Pups were then collected at P0. We analyzed the kidney cortex and observed a strong streptavidin signal within the glomeruli of P0, *Nphs2*^*BioID2/+*^ mice (Fig 3D, arrowheads). Contrarily, we did not observe signal in wildtype control littermate mice. Together, these data highlight the efficacy and specificity of our podocin-BioID model which can be utilized across a spectrum of ages.

**Figure 3.**
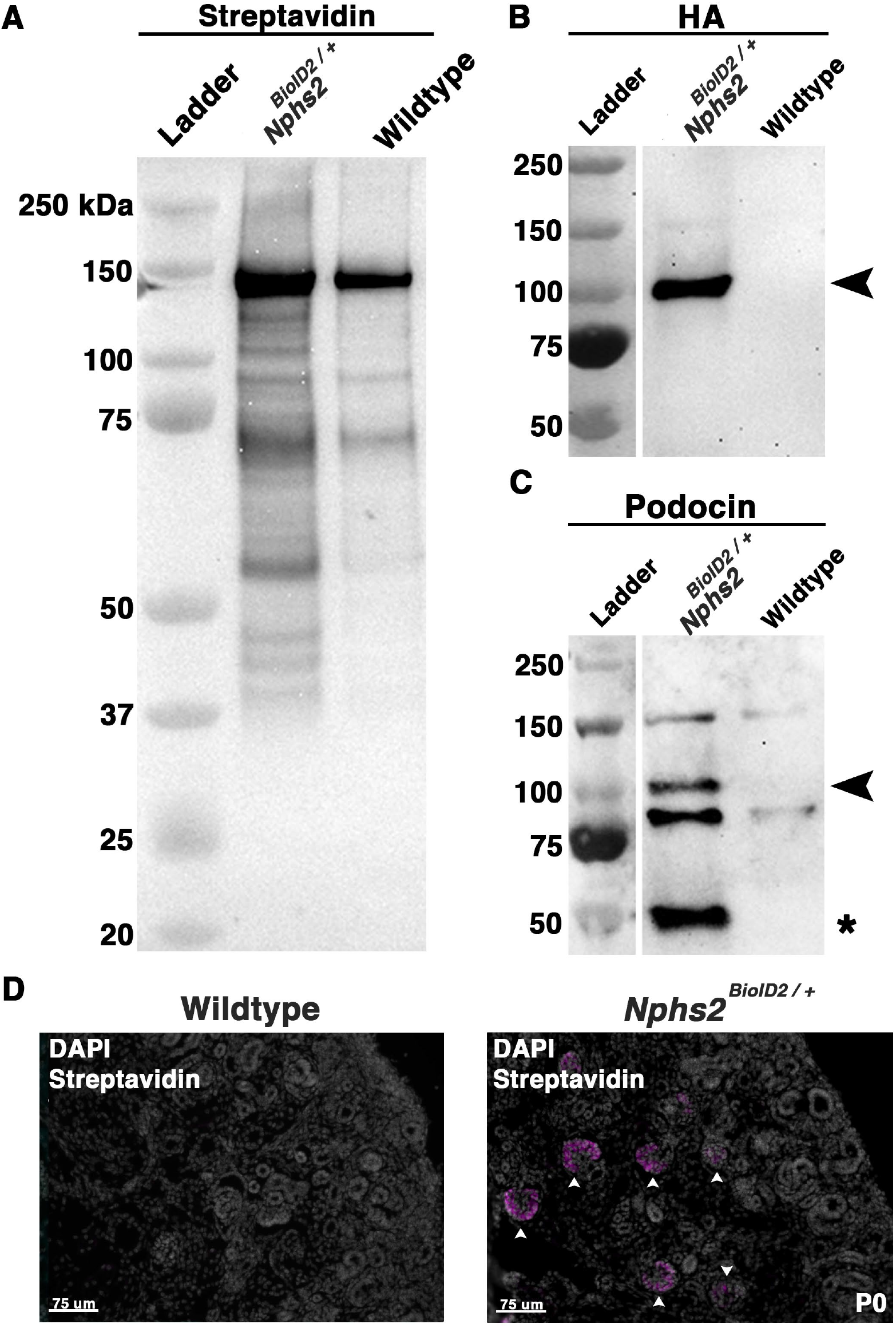
Biotin administered *Nphs2*^*BioID2/+*^ mice show an enrichment of biotinylated proteins specifically within glomeruli. **(A)** Streptavidin bead purified lysates from biotin injected *Nphs2*^*BioID2/+*^ and wildtype age matched 8–10-week-old littermate controls were subjected to protein separation, Western blot, and probed for streptavidin. **(B)** Lysates blotted with an anti-HA antibody show robust and specific detection of the HA signal only within the *Nphs2*^*BioID2/+*^ sample (arrowhead). **(C)** Samples probed for podocin show two specific bands for both BioID2-tagged podocin (arrowhead) and endogenous podocin (asterisk). **(D)** Micrographs of P0 kidney sections showing DAPI (grey) and streptavidin (magenta) labeling. Pregnant dams were injected with 5 mg/kg of biotin from E11.5 to E18.5. Arrowheads show glomeruli labeled with streptavidin. Scale bar: 75 μm.

### Identification of the podocyte foot process proteome by mass spectrometry profiling

Kidney cortices of three 8 to 10-week-old, sex matched mice were collected as one biological replicate, after biotin administration for 1 week. All MS analyses were run in triplicate *i*.*e*. 9 mice per condition, totaling 18 mice (*Nphs2*^*BioID2/+*^ vs control littermates) per MS analysis. To representatively capture the podocyte foot process proteome, we combined the results of three separate MS analyses (Figure 4). Significant sex differences were not apparent in our studies (Supplemental Tables II-IV). From our proteomics analysis 11 proteins were found across all three MS analyses to have an averaged log_2_ fold change ≥ 2.5. Many of these top proteins including podocin, Kirrel, Tjp1, Pard3b, Magi2, Dnd, and Synpo are documented to localize to the slit diaphragm.^37^ Yet others including an Immunoglobulin like domain containing receptor 2 (Ildr2) and a Formin binding protein 1 like (Fnbp1l, also known as Toca1) were, until now, unreported components of the podocyte foot process/slit diaphragm. Additionally, 6 proteins were identified across two MS analyses as significantly enriched with an average log_2_ fold change ≥1.75. These include documented slit diaphragm components Tjp2 and Cd2ap (Figure 4A). All top 17 proteins had a stringent log_2_ fold change ≥ 1.75. Tables listing the significant foot process proteins identified from each MS analysis are reported in Supplemental Tables II – IV. To add support to our relative log_2_ fold cut off, we assayed an immunoglobulin superfamily adhesion molecule, Jam1/F11r,^38^ with an observed log_2_ fold expression change of 0.7 (Supplemental Table II). We identified Jam1 expression in cells of the proximal convoluted tubule, abutting the glomerulus, (Supplemental Figure 1) but not within the glomerulus. Potentially, non-glomerular cortex proteins that are biotinylated are also isolated although these appear minimal in our findings. However, this helped establish a relative fold-cutoff for which we start to identify non-podocyte proteins.

**Figure 4.**
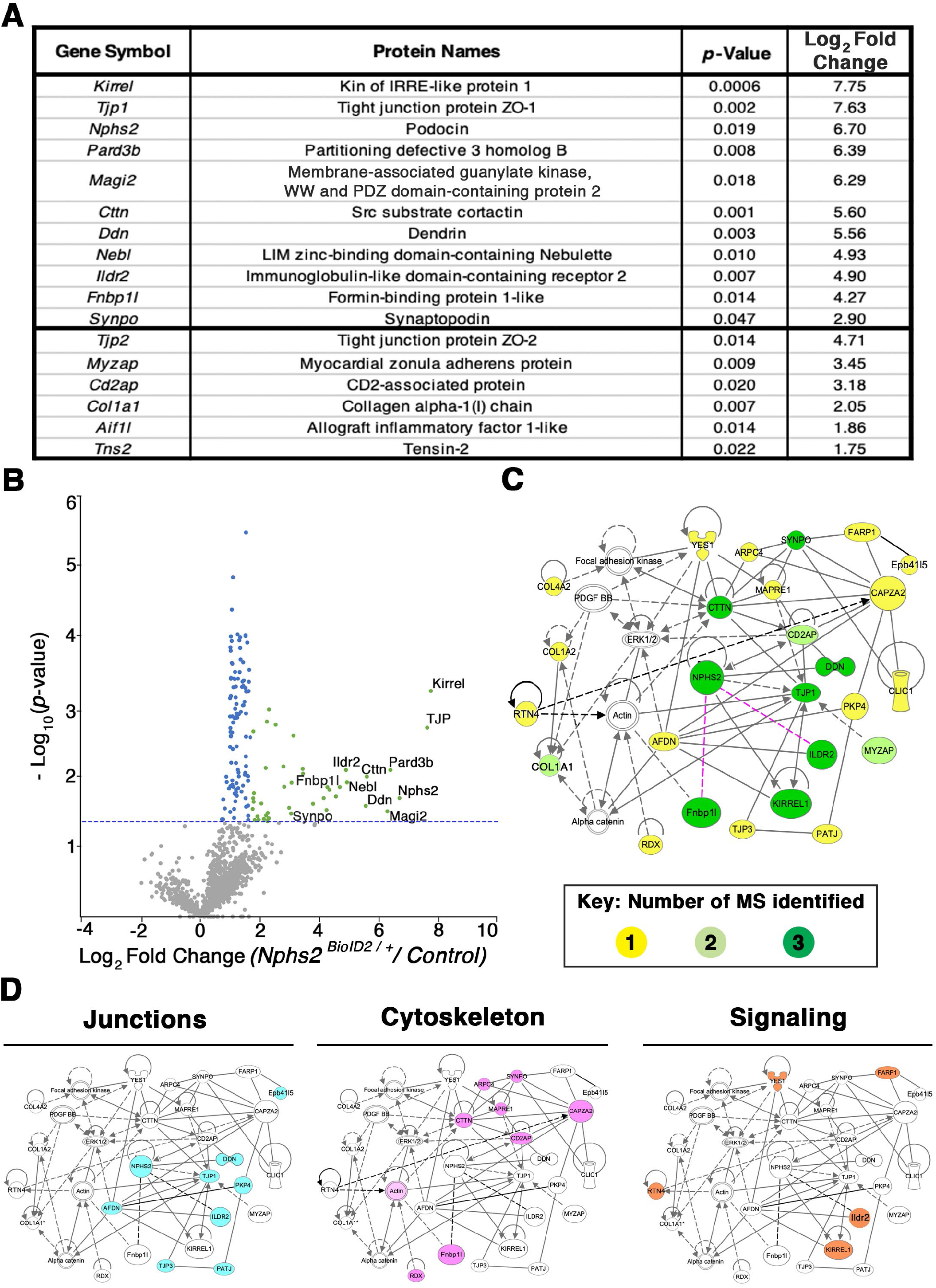
Proteomics profiling of the podocyte foot process identifies documented slit diaphragm components and novel candidates with unexplored podocyte function. **(A)** Three separate MS analyses were combined, and the average log_2_ fold change and respective *p*-values were averaged to produce a list of the top foot process proteins. The top 11 proteins were identified across all three MS profiles. Six additional proteins were identified across two of the three MS analyses, denoted following the thick black bar. **(B)** A volcano plot depicting approximately 1400 proteins identified across all three MS analyses. Green dots represent the 40 proteins identified across three separate MS profiles as having a log_2_ fold change ≥ 1.7, and *p*-value ≤ 0.05, blue dots denote all proteins identified to have a significant *p*-value ≤ 0.05, and grey dots are proteins with a *p*-value ≥ 0.05. The top 11 proteins consistently uncovered across all MS analyses are embedded with gene symbols in the plot. The blue dotted line represents a *p*-value ≤ 0.05. **(C)** Network topology was generated utilizing *Qiagen Ingenuity Pathway Analysis* (IPA) on the top 54 proteins from our cumulative proteomic profiles. IPA produces a proposed web of relationships from published literature with most of our top proteins represented, (11 of 17), with documented connections to other foot process and slit diaphragm components. The color intensities of each protein (yellow to green) represent the number of MS analyses from which each protein was identified. Darkest green shade=all three MS analyses, lighter green shade=two of three MS analyses, and yellow=a single MS analysis. Dotted lines and unshaded proteins represent predicted interactions/interactors from IPA analysis, respectively. Magenta lines are novel proteins identified in this study to be present within the podocyte foot process. **(D)** Dissecting the IPA network, identifies three central nodes representing proteins associated with junctions (cyan), cytoskeleton (magenta), and signaling (orange).

To surmount a complete list of podocyte foot process proteins we established a log_2_ fold change cut off at 1.2 across all three mass spec analyses, and excluded histone, ribosome, and mitochondrial proteins to arrive at a catalog of 54 proteins (Supplemental Table V). Graphically depicting the compiled proteome in a volcano plot with the total ∼1400 proteins identified and cataloging proteins with a log_2_ fold change ≥ 1.75 and *p*-value ≤ 0.05 in green, proteins found to have a *p*-value ≤ 0.05 in blue and proteins not found to be significant in grey (Figure 4B, Supplemental Table V). The blue dotted line represents a -log_10_ (*p*-value≤ 0.05) (Figure 4B). All significantly identified proteins (*p* ≤ 0.05) had at least two unique peptides identified via MS analysis. The most highly detected and significant proteins from our proteomic profiling, clustering with documented slit diaphragm components, are found in the right scatter of the volcano plot, depicted with green dots (Figure 4B, Supplemental Table V).

To interrogate our proteomic findings further, the top 54 podocyte foot process proteins were utilized for *in silico* analyses. We input Supplemental Table V into *Qiagen Ingenuity Pathway Analysis (*IPA) and performed a variant effect analysis to compute a proposed interactome based on published literature (Figure 4C). The representative web of interactions from IPA was color coded based on the number of MS analyses each protein appeared in, one (yellow), two (light green), and three (dark green) (Figure 4C). Two novel foot process proteins were identified by IPA, Ildr2 and Fnbp1l, to be involved in the interactome with connections to Afadin and actin, respectively (Figure 4C). We added two dotted magenta lines for Ildr2 and Fnbp1l potential interactions with podocin (Figure 4C). Unshaded proteins and dashed lines are predicted interactors/interactions from IPA. Our IPA proposed network includes 26 of our 54 input foot process proteins. The IPA interactome identifies three main nodes, *i*.*e*. junctions, cytoskeleton, and signaling (Figure 4D). We highlighted the respective proteins that contribute to each node for junctions (cyan), cytoskeleton (magenta), and signaling (orange) (Figure 4D). We next input these 54 proteins into the *Database for Annotation, Visualization, and Integrated Discovery (DAVID)* to identify the gene ontogeny (GO) terms that are most highly enriched across our foot process proteome. DAVID analysis identified cytoskeleton protein binding, cell-cell junctions, and actin filament-based processes as the top molecular function, cellular component, and biological processes GO terms, respectively (Figure 5A). Additionally, DAVID and IPA overlap in their representation of junctional signaling and actin binding as top GO terms and canonical pathway, respectively. Taken together, these findings align with the major functional roles of podocin and other foot process/slit diaphragm components.

**Figure 5.**
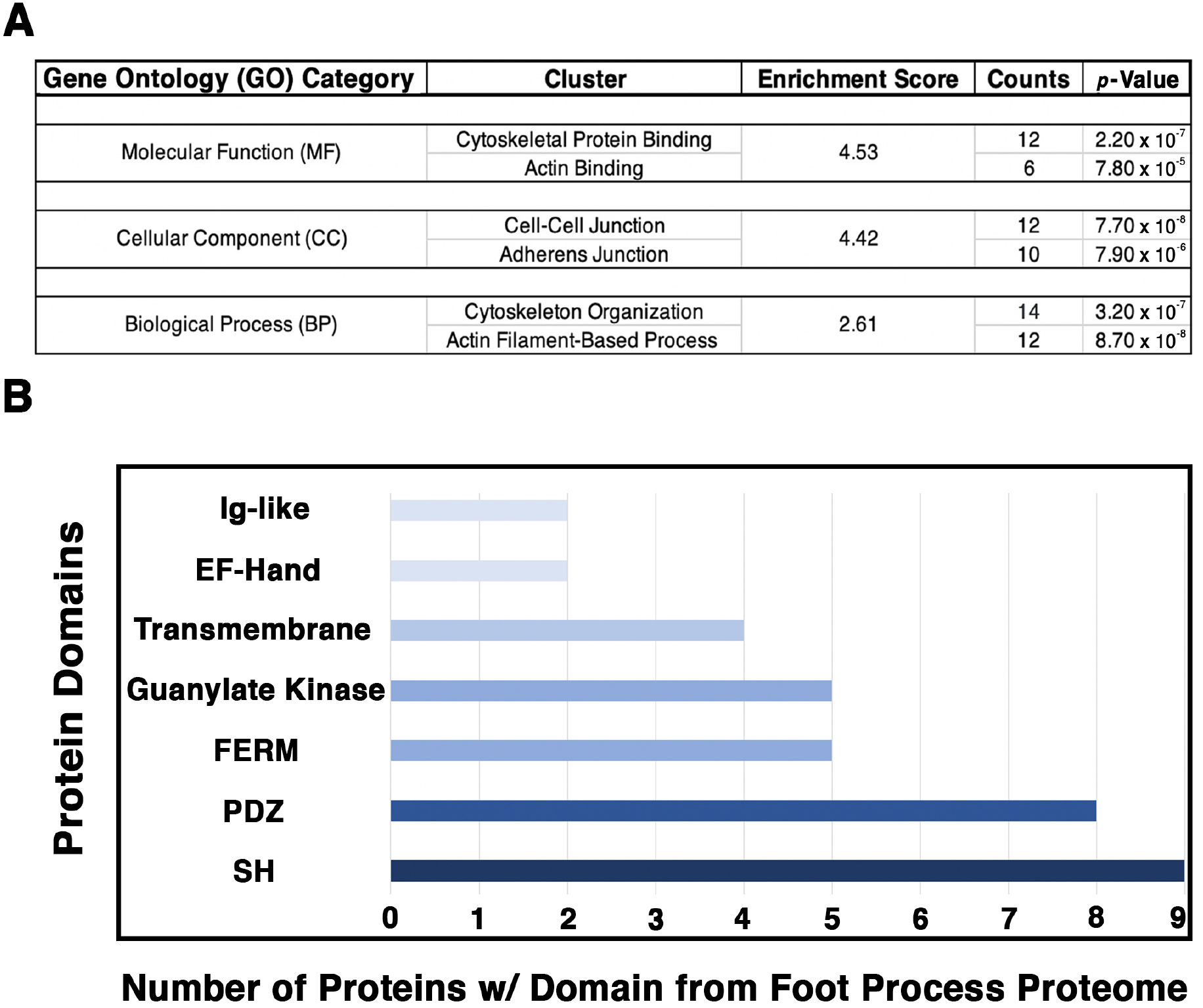
Analysis of the podocyte foot process proteome identifies the cytoskeleton, cell-cell junctions, and actin-based processes as the top GO categories with respective protein domains that align with these functions. The Database for Annotation, Visualization, and Integrative Discovery (DAVID) was utilized for GO clustering analysis of the top 54 proteins identified to have a log_2_ fold change ≥ 1.20, and *p*-value ≤ 0.05 (Supplemental Table V). **(A)** Overlap in GO readout was observed across the three separate GO categories analyzed, molecular function, cellular component, and biological process, with cytoskeleton, actin, and cellular junctions being the top hits. The respective top clusters within each GO category are listed, alongside the relative enrichment score, number of proteins, and *p*-values. **(B)** Protein domain analysis of the top 54 podocyte foot process proteins provides evidence for protein-protein interactions and likely scaffold and protein-protein complex formation. Each protein’s respective protein domains were binary counted for presence or absence within the proteomics profile. The top seven protein domains identified from the podocyte foot process proteome consist of a Src homology (SH) (n = 9 proteins), PSD-95, Disc large, and ZO – 1 (PDZ) (n = 8), 4.1 protein, Ezrin, Radixin and Moesin (FERM) (n = 5), Guanylate Kinase (n = 5), Transmembrane (n = 4), E and F helix – Hand (EF – Hand) (n = 2), and an Immunoglobulin - like (Ig – like) domain (n = 2). Cumulatively, these 7 domains represent ∼ 50 % of the top proteins identified from the foot process proteome (26 / 54).

To further decipher the contributions of our top 54 podocyte foot process proteins to junctions and cell-cell contacts we cataloged the protein domains represented within each protein. We utilized binary counting for the presence and absence of a domain. We did not account for the number of same/similar domain(s), *i*.*e*. SH2/3, within a protein. We identified 7 top protein domains within the catalog of the podocyte foot process proteome that included SH, PDZ, FERM and Ig-like domains among others (Figure 5B). All of these top protein domains align with the nodes of the podocyte foot process proteome, *i*.*e*. junctions, cytoskeleton, and signaling.

### Ildr2 and Fnbp1l are expressed by podocytes and co-localize with podocin

Our proteomic analyses uncovered several proteins which have not been previously described to localized to the podocyte foot process or have podocyte-specific functions. We therefore wanted to validate their expression in podocytes and any colocalization with podocin. We utilized immunofluorescence on kidney sections to visualize the location of three proteins in particular: Myozap, Fnbp1l, and Ildr2. First, a known slit diaphragm and foot process component which was identified in our proteomics, Pard3b, was assessed for its colocalization with podocin. Pard3b both exhibited positive staining in the glomerulus and overlapped with the HA signal of podocin-BioID (Figure 6A). Pard3b positive staining is also found in the adjacent tubule cells where it helps maintain epithelial integrity (Figure 6A).^39^ Myozap regulates cardiac function through Rho-dependent activity and interacts with junctional proteins such as ZO-1^40^. It was identified across two of the three MS analyses (Figure 4A) and therefore we decided to investigate its localization. Utilizing an antibody against a synthetic human MYOZAP peptide,^41^ we identified strong signal of Myozap in the kidney endothelium, both in the glomerular capillaries and outside of glomerular structures (Supplemental Figure 2). Myozap exhibits some degree of overlap with podocin, however, the majority of Myozap protein detection does not colocalize with podocin, suggestive of additional roles outside of podocytes (Supplemental Figure 2).

**Figure 6.**
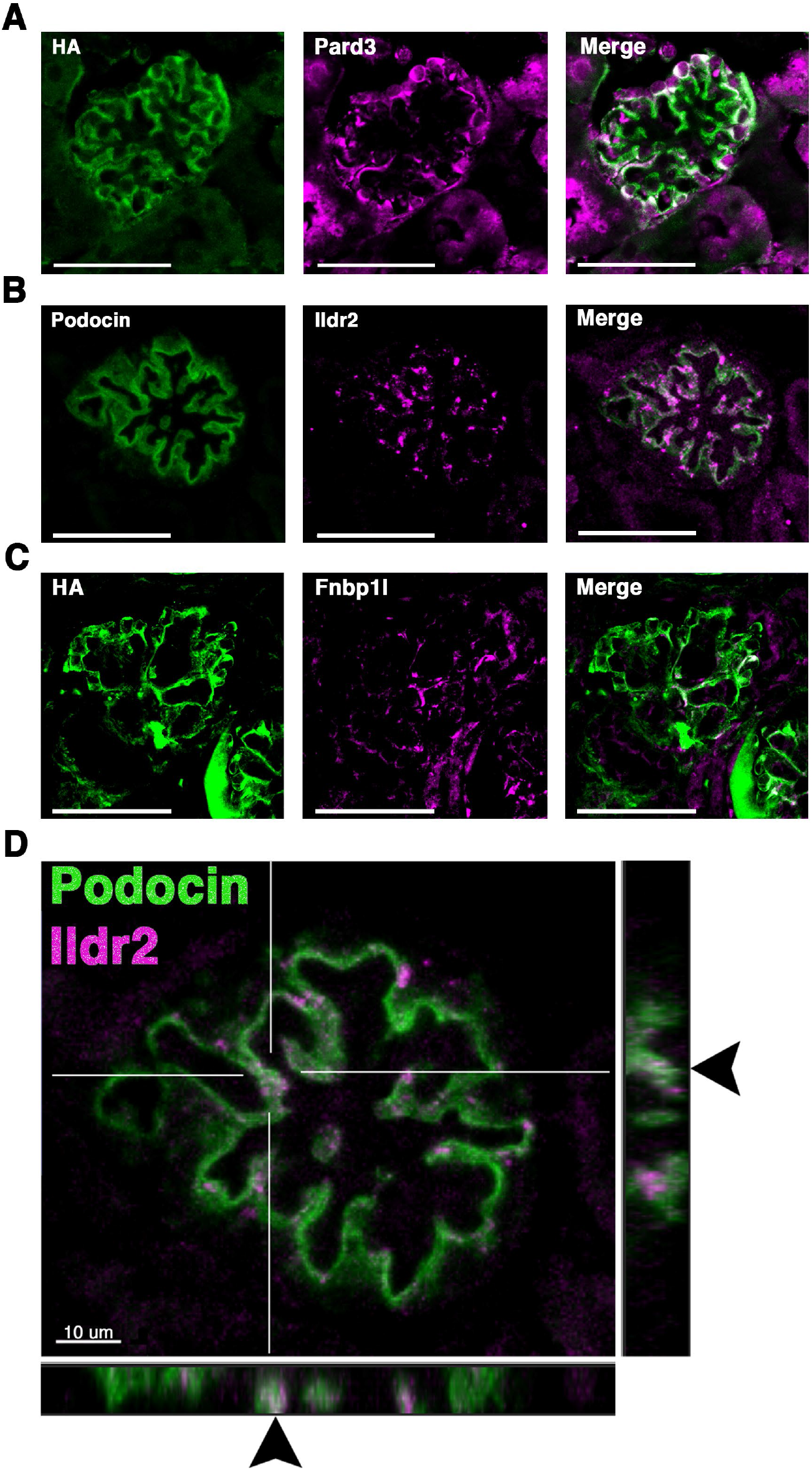
Two novel foot process candidates, Ildr2 and Fnbp1l, localize to podocytes and overlap with podocin. **(A)** Micrographs showing Pard3 (magenta) localization within podocytes and colocalization with the HA signal of podocin-BioID (green). **(B)** Co-immunostaining of Ildr2 (magenta) and podocin (green) within the glomerulus. **(C)** Fnbp1l (magenta) co-localization with the HA signal of podocin-BioID (green) in glomeruli. Scale bars in A-C: 50 μm **(D)** A merged image of podocin (green) and Ildr2 (magenta) through a confocal z-projection shows punctate localization of Ildr2 and overlap (arrowheads) with podocin in the z-plane in the left and bottom panels. Scale bar in D: 10 μm.

We identified Fnbp1l, a documented junctional and actin organizing protein,^42, 43^ across all three MS analyses with a log_2_ fold change ≥ 4 (Figure 4). Fnbp1l has been identified in human and mouse podocytes from single cell RNA-seq analysis and linked to podocyte cytoskeleton dynamics *in vitro*, although it has not been documented to colocalize with the slit diaphragm or foot process-associated proteins.^44,45^ We found that Fnbp1l co-localizes with podocin in continuous stretches within glomeruli observed as a white signal from the Fnbp1l (magenta) and podocin (green) overlap (Figure 6C). *In situ* hybridization confirms the glomerular expression of *Fnpb1l* (Supplemental Figure 3C’ arrowhead), similar to *Nphs2* (Supplemental Figure 3A), in addition to a tubular expression pattern at postnatal day 2 (P2).

Ildr2, a member of the B7 superfamily of immunoglobulins, is found highly enriched in the podocyte foot process proteome across all MS profiles with an average log_2_ fold change of 4.9 (Figure 4). Ildr2 is a member of the angulin family and localizes to tricellular junctions of *in vitro* cultured epithelial cells^46^. In the kidney, Ildr2 exhibited a punctate staining pattern that colocalized with podocin in adult mouse glomeruli, reminiscent of the punctate and restricted pattern of tricellular junction staining observed *in vitro* (Figure 6B, 6D arrowheads)^46^. In the z dimension we observe overlap of Ildr2 (magenta) with podocin (green) as white punctate foci denoted by arrowheads (Figure 6D). We further validated *Ildr2* expression within glomeruli via *in situ* hybridization. *Ildr2* expression is found in glomeruli (Supplemental Figure 3B’, arrowhead) and tubule cells of the kidney (Supplemental Figure 3B’, arrow) in P2 mice. We went on to verify Ildr2 localization within early renal vesicles, comma, and S-shaped bodies of developing nephrons at E15.5 (Figure 7B, Supplemental Figure 4). Ildr2 is more membranous and contiguous yet with some punctate detection at E15.5 both within podocytes and cells of the developing nephron tubule (Figure 7B). Collectively these studies provide evidence that Ildr2 and Fnbp1l are novel podocyte foot process and potentially slit diaphragm components that likely play roles in helping to maintain podocyte architecture.

**Figure 7.**
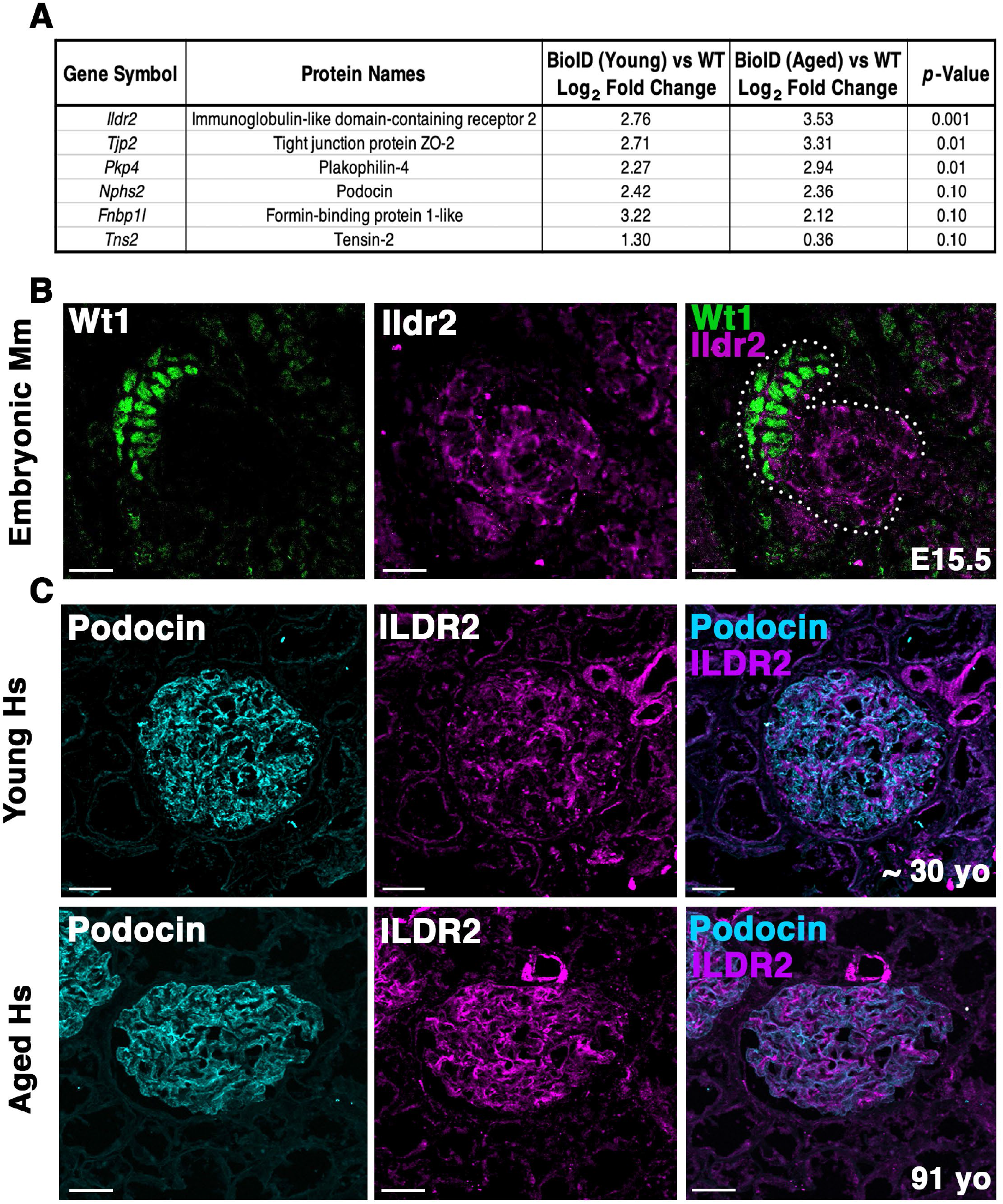
Ildr2 is detected in developing nephron structures and shows increased signal in aged mouse and human glomeruli compared to young. **(A)** Proteomic profiling of two aged (∼2 years; 108 week old) male *Nphs2*^*BioID2/+*^ mice compared to 8-10 week old *Nphs2*^*BioID2/+*^ mice. Table lists 3 significantly increased proteins, podocin, Fnbp1l, and one downregulated protein. All proteins were detected with at least 2 razor unique peptides. **(B)** Micrographs of E15.5 kidney sections showing Ildr2 (magenta) in developing nephron structures (dotted outline) of mice (Mm; Mus musculus) with podocytes denoted by the Wt1 (green). Scale bar: 20 μm. **(C)** Micrographs of sections from young (∼30yo) and aged (91yo) human (Hs; Homo sapiens) kidneys tissue co-immunostained for ILDR2 (magenta) and podocin (cyan). Scale bar: 50 μm.

### Ildr2 protein levels increase in both mouse and human glomeruli with age

To determine whether our podocin-BioID model can detect changes to the localized proteome in aging, we performed proteomics on ∼2-year-old (108 weeks) male *Nphs2*^*BioID2/+*^ mice and compared the results to our 8–10-week-old *Nphs2*^*BioID2/+*^ male mice to assess changes to the proteome (Figure 7A). We identified significant increases in Tjp2 and Pkp4 and the largest change (2.76-fold) in Ildr2. We were unable to detect any significant decrease in proteins at *p* ≤ 0.05, although Fnbp1l and Tns2 showed a decreasing trend. We observed minimal change in podocin when comparing the two age groups (Figure 7A). To assess the human relevance of our findings, we assayed ILDR2 immunofluorescent staining in young (age ∼30 years) and aged (91-year-old) human kidney sections. ILDR2 displayed a similar punctate staining pattern in young human glomeruli. In correlation with our findings from the mouse, we found an increase in ILDR2 staining in the aged glomeruli. Interestingly, the expression pattern no longer displays a punctate pattern but rather a more diffuse membranous staining pattern, similar to the mouse E15.5 stage (Figure 7B,C). To verify the specificity of the increased ILDR2 signal to podocytes, we co-immunostained kidney sections with ITGA8 (mesangial marker) and CD31 (endothelial marker) (Supplemental Figure 5). We found minimal overlap of ILDR2 with either ITGA8 or CD31 and the most significant overlap of signal with podocin, albeit more diffuse (Figure 7C, Supplemental Figure 5). We quantified the corrected total glomerular florescence (CTGF) by selecting the glomerulus as a region of interest (gROI) then quantifying total florescence intensity and subtracting out the mean background florescence for the glomerular area, *i*.*e*. total gROI florescence – (glomerular area x mean background florescence) = CTGF, (Figure 8). We confirmed a significant (*p* ≤ 0.007) increase in ILDR2 in aged human glomeruli compared to young human kidney tissue (Figure 8) validating the ability of our podocin-BioID mouse model to detect conserved changes in proteomes with aging.

**Figure 8.**
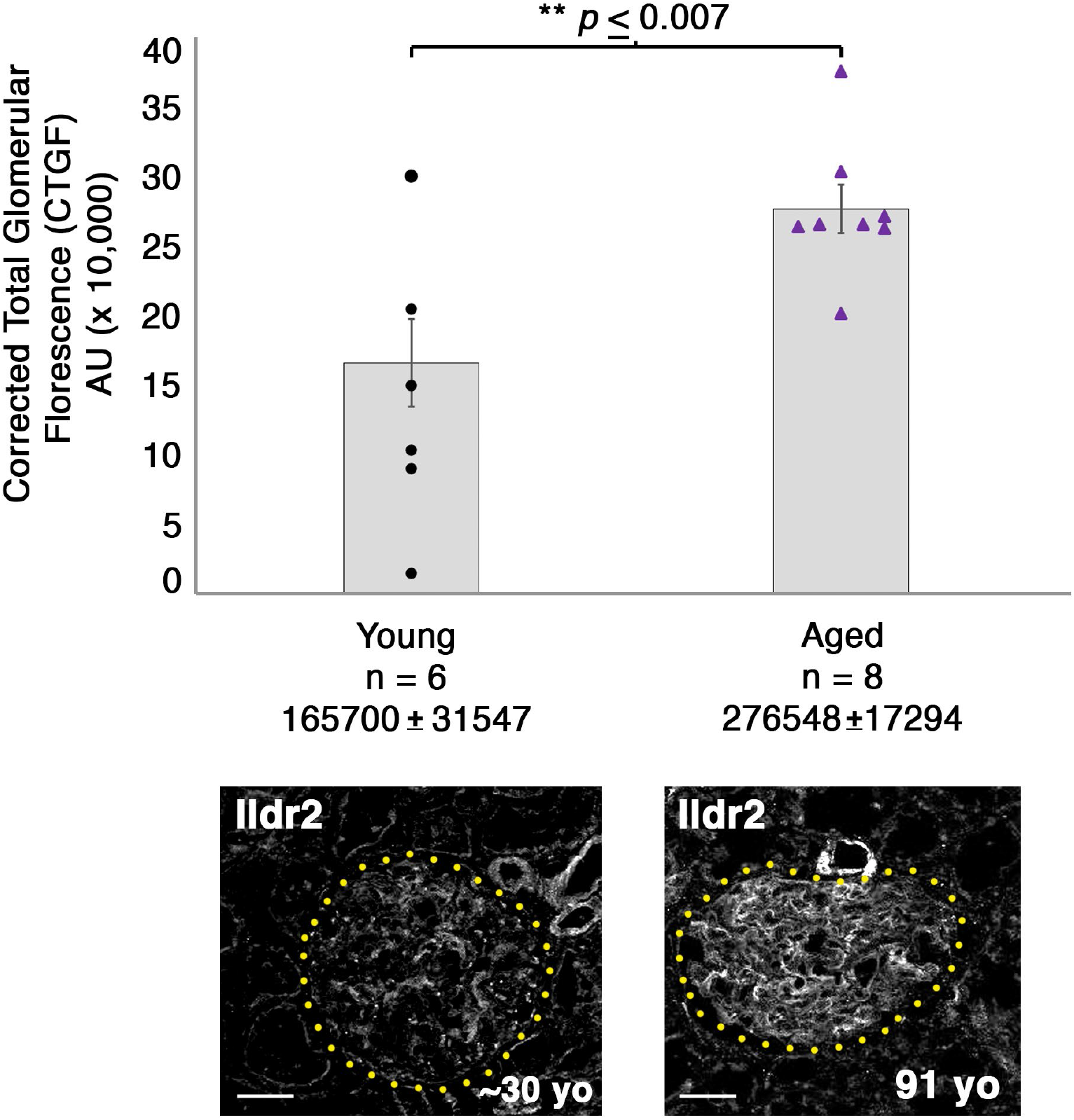
Aged glomeruli exhibit a significant increase in ILDR2 corrected total glomerular florecence (CTGF) compared to young human glomeruli. ILDR2 CTGF was calculated by selecting the gomeruli as the ROI then measuring the total florescence within the area and subtracting out the sum of the area by the average background florecence, *i*.*e*. Total gROI florescence – (Area of glomerulus *x* mean of background florescence) = CTGF. Two example glomeruli are shown from each age and outlined in yellow dots to depict the respective ROI utilized to calculate CTGF. Scale bar: 50 μm.

## Discussion

Podocytes are extraordinary epithelial cells of the kidney that intertwine their foot process extensions to establish a cellular junction, the slit diaphragm, that is distinct from other cellular junctions in the human body. The slit diaphragm was identified more than five decades ago as an electron dense region between two podocyte foot processes, as visualized by EM.^47,48^ Many proteins that compose the slit diaphragm when mutated are associated with nephrotic diseases, including nephrin, podocin, Magi2, and Cd2ap.^49^ Loss of podocyte integrity is one of the most common clinical observations in kidney disease. End stage renal disease being a top 10 cause of death in the US necessitates the need to identify novel components of the slit diaphragm, and how the slit diaphragm changes with disease, to help spur the development of new therapeutic options and identify biomarkers of disease severity.

We utilized a new *in vivo* biochemical tool to interrogate the proteome of the podocyte foot process *via* knock-in of a BioID moiety, generating our podocin-BioID model. The sensitivity and advantage of BioID is that it allows for weak and transient interactions to be identified, in addition to withstanding harsh isolation conditions.^6^ Further, the biotin-streptavidin bond is one of the strongest known non-covalent interactions, enhancing the isolation of biotinylated proteins via streptavidin-coated magnetic bead. We validated our model recapitulates normal podocin localization by TEM and showing podocin-BioID localizes normally to the slit diaphragm. Activity of the biotin ligase was confirmed through probing kidney sections and lysates with streptavidin and finding an enrichment of biotinylated proteins. Our proteomics profiling identified more than 50 candidate molecules with significant enrichment in the podocyte foot process, log_2_ ≥ 1.2. Within this dataset we were able to identify protein interactors known to associate with podocin as well as novel candidates not found in previous investigations of the slit diaphragm and the podocin-associated proteome^22,23^. Additionally, we were able to identify changes to the localized proteome that occur with aging. It would be interesting to compare the proteome across additional stages such as during development. Podocytes begin as a columnar epithelium with only tight and adherens junctions which remodel and later mature to form the specialized slit diaphragm complex.^16^ With the ability of biotin to cross the placental barrier, the proteome could be compared from development to maturity to identify changes that may help inform how these specialized junctions are formed.

The proteomics analysis revealed potential nuances of the podocyte foot process that are still under investigation. We identified 11 proteins that repeated over all three MS profiles. However, there were more than 30 proteins that only surfaced in a single MS analysis and split across the three MS analyses. While this may be due to experimental variability, such as in the isolation of proteins, this may also highlight some differences in biological activity in the foot process. The dynamics, variability, and protein turnover in the podocyte slit diaphragm is relatively unknown. Pointing to the relative variability of one documented component, Synaptopodin (Synpo), across the three MS analyses the respective log_2_ fold changes were observed at 1.0, 2.9, and 4.8 which indicates potential protein turn over, variability in the slit diaphragm components, or potential variability of the MS detection. Further still, a second documented slit diaphragm protein, Cd2ap, was only identified in two of the MS profiles. These cofounding issues made a single MS analysis a limited view, or snapshot, of the proteins present within the podocyte foot process. By combining three separate MS analysis we have uncovered a more complete profile of the podocyte foot process and slit diaphragm. Many of the top candidates identified in the podocyte foot process proteome are well documented slit diaphragm components including Kirrel, Nphs2, Par3, Magi1/2, Tjp1/2/3, Dnd, Synpo, and Cd2ap. Furthermore, our *in silico* analysis utilizing DAVID highlights anticipated GO terms, including actin binding, cell-cell junctions, adherens junctions, and cytoskeleton organization that align with podocyte function. We conclude the podocin-BioID model provides a spatial-specific approach to identify proteins that are notoriously difficult to isolate.

We cannot rule out that biotin delivery, metabolism, and selective labeling may limit some proteins from being detected. One report depicts a time lapse of biotin administration *in vivo* with positive streptavidin enrichment for as long as 18-hour after biotin administration.^50,51^ However, biotin requires an accessible primary amine within a peptide to biotinylate. Proteins with little open structure or a limited number of lysine residues could be missed by the labeling strategy. The possibility that proteins were missed is brought to light by the inability to identify nephrin (*Nphs1*) across any of the mass spec analyses run. One rational for the inability to identify nephrin, a known slit diaphragm molecule that has been shown to interact with podocin, posits that its C-terminal domain is only minimally assessable to podocin, while the remaining majority of the 180 kDa protein is spanning the extracellular space of the slit diaphragm.^27^ Therefore, while there are significant advantages to the BioID system over traditional immunoprecipitations followed by MS, it is also subject to missing important interactions. Additionally, we did not identify a significant enrichment of solute transporters as in a recent MS analysis of the podocin-associated proteome.^23^ This may represent a more restricted localization of the modified podocin-BioID protein along the foot process, the inability to biotinylate these proteins, or the lack of enrichment in our studies due to their normal biotinylation without the addition of excess biotin. Together, these findings highlight the efficacy of using or combining multiple approaches to obtain a more complete understanding of the localized proteome.

One of the novel foot process candidates we identified, Fnbp1l, has multiple roles in microtubule binding, cell polarity, motility, actin organization, junctional localization and signaling.^42,43^ Based on these roles, we hypothesize that Fnbp1l similarly helps maintain the slit diaphragm and foot process integrity through associations with the actin cytoskeleton and links to the slit diaphragm. Single cell RNA-seq analyses have previously found *Fnbp1l* expression in human^44^ and mouse podocytes and tubule cells.^45^ From the Kidney Interactive Transcriptomics (KIT), *Fnbp1l* is most highly expressed in podocytes compared to the tubule cells of the human kidney.^52^ Further, *Fnbp1l* is also expressed in the zebrafish pronephros^53^ with a similar pattern to other zebrafish tight junction proteins such as ZO-1/2^54^, however the specific expression of *Fnbp1l* within the zebrafish podocytes remains unknown. With the conservation of *Fnbpl1* expression in the renal system across vertebrates, it likely plays an important role in helping maintain epithelial integrity in the various cells of the nephron, including podocytes.

Ildr2 was identified in the podocyte foot process/slit diaphragm proteome across all three MS profiles with a log_2_ fold change ≥ 4.5 and a *p*-value ≤ 0.007. Ildr2 belongs to the B7/CD28 family of proteins, encompassed by the Ig superfamily (IgSF), with pivotal roles in immunomodulation and maintenance of peripheral self-tolerance.^55^ Ildr2 evinces immunomodulatory activity, wherein administration of Ildr2 as a fusion protein (ILDR2-Fc) rebalances immune homeostasis, which leads to an amelioration of autoimmune disease states in mouse models of rheumatoid arthritis, type I diabetes, and relapsing-remitting multiple sclerosis.^55-56,57^ Recently, an Ildr2 blocking antibody, BAY 1905254, was generated to block the immunosuppressive activity of Ildr2 for cancer immunotherapy.^57^ BAY 1905254 promotes T-cell activation *in vitro* and enhances antigen-specific T-cell proliferation and cytotoxicity *in vivo*, and is currently in phase I clinical trials.^57^ Ildr2 presents an unexplored niche in podocyte biology centered around the function of podocytes in immune cell modulation. Yet, the ability for immune cells such as T-cells to interreact with podocytes is challenged by the podocyte cellular environment and the GBM. Therefore, whether Ildr2 could play an immune related role in podocytes remains tentative.

*Ildr2/ILDR2* is found to be specifically expressed in mouse and human podocytes from publicly available databases including KIT.^44,58^ *Ildr2* was also identified as a Wt1 transcriptional target in podocytes by ChIP-seq.^59^ In early embryonic and postnatal mouse kidneys, *Ildr2/*Ildr2 is expressed and localized to tubule cells and podocytes. However, in the adult kidney Ildr2 localization becomes restricted to podocytes. At the timepoints of podocyte development, we observe a more membranous and contiguous pattern of Ildr2 in comma and S-shaped bodies, with some foci of heightened detection. In mature glomeruli, Ildr2 is restricted to punctate foci that colocalize with podocin. Ildr2 has been identified at the site of tricellular tight junctions (tTJ) in murine retinal pigment epithelium and *in vitro* within EpH4, a mouse mammary gland cell line.^46,56^ Additionally, Ildr2 has recently been identified to interact with Afadin in human embryonic kidney cells (HEK293).^60^ Afadin is also reported to localize to the site of specialized tricellular junctions and mediate mechanotransduction.^61^ The role of mechanical strain present from the underlying fluid flow sheer stress on these specialized tTjs and recruitment of specific proteins to these domains remains unresolved. The function and even the localization of tTJs in podocytes is poorly understood. Many proteins identified in the podocyte foot process proteome are documented junctional proteins containing classified SH, PDZ, FERM, transmembrane, EF-Hand, and Ig-like domains. Yet, Ig-like domain containing proteins were a considerably smaller population. It is intriguing to speculate that Ig-like domains or a combination of PDZ, SH, and Ig-like domain aids in the recruitment of specific proteins to specialized junctions such as tTJs and the slit diaphragm. Further still, the requirement of these proteins for managing stress such as from changes in fluid flow from the fenestrated endothelium and conversely the potential requirement of mechanostress for recruitment of proteins to these tTJs and slit diaphragms remains an area in need of further investigation.

In our aged podocin-BioID mice as well as our aged human kidney sample, we identified an increase in Ildr2 within podocytes. The pattern of Ildr2 is more membranous with some punctate foci in embryonic renal structures yet becomes detectable only as punctate foci as the mouse matures. Proteomic profiling of kidneys from ∼2-year-old mice showed a significant enrichment and increase of Ildr2 versus 8-10-week-old kidneys. We validated ILDR2 significantly increases in aged (91yo) human glomeruli via quantifying the corrected total glomerular florescence of young and aged human glomeruli. We further noted that in the aged human glomeruli ILDR2 signal is more abundant in zones where podocin signal is less and conversely where podocin signal is high ILDR2 is less abundant, although there is still overlap between the two proteins. The ILDR2 signal appears to be specific to podocytes, as minimal overlap of ILDR2 signal was detected with mesangial (ITGA8) and endothelial (CD31) markers. Aged human glomeruli present an ILDR2 pattern that more closely resembles the mouse embryonic stage with a contiguous pattern rather than the punctate pattern of 8–10-week-old mice. We hypothesize that Ildr2 is a tight junction component and increasing Ildr2/ILDR2 protein in aged podocytes helps maintain podocyte integrity. This likely allows podocytes to withstand physiological changes that occur with aging through the modification of junctional components. The additional increase of tight junction and desmosome associated proteins Tjp2 and Pkp4, respectively, in our aged mouse proteome further supports this hypothesis.

In this study, we developed an innovative tool for the identification of novel proteins within spatially restricted podocyte foot process/ slit diaphragm. Current efforts are now aimed at uncovering proteins differentially regulated during slit diaphragm development, aging, and in renal disease. It is crucial we uncover the changing composition of the podocyte slit diaphragm and foot process as it holds potential for new therapeutic treatment targets to either preserve or prevent the loss of podocyte integrity in kidney disease.

## Methods

### Genetic CRISPR/Cas9 engineering of the Nphs2 locus to append a BioID2 moiety

CRISPR/Cas9 targeting, and donor vector design were performed by UNC Chapel Hill Animal Models Core. *Benchling* software was used to identify Cas9 short guide RNAs (sgRNA) overlapping the podocin (*Nphs2*) stop codon. Guide RNAs were cloned into a T7 promoter vector followed by *in vitro* transcription and spin column purification. Functional testing was performed by transfecting a mouse embryonic fibroblast cell line with sgRNA and Cas9 protein (produced and purified by the UNC Protein Expression Core). The sgRNA target region was amplified from transfected cells and analyzed by T7 endo1 nuclease assay (New England Biolabs). sgRNA, *Nphs2*-g78B (protospacer sequence 5’-gCCATTCGCCTATAACAT-3’; lower case g indicates heterologous guanine added at 5’ end of native sequence for efficient T7 *in vitro* transcription) was selected for genome editing in embryos. *Nphs2* was amplified from adult mouse kidney cDNA and cloned into the MCS-13X Linker-BioID2-HA plasmid (MCS-13x Linker-BioID2-HA was a gift from Kyle Roux (Addgene plasmid # 80899 ; http://n2t.net/addgene:80899 ; RRID:Addgene_80899).^30^ A donor vector was subsequently constructed from the *Nphs2*-13x Linker-BioID2-HA plasmid with the following features: (1) a 1018 bp 5’ homology arm encompassing sequence immediately 5’ of the *Nphs2* stop codon including 276 bp coding sequence from *Nphs2* exon 8, (2) a 267 bp in-frame Glycine/Serine-rich linker sequence, (3) a 696 bp coding sequence for the humanized biotin ligase of *A. aeolicus* with a R40G mutation in the catalytic domain (BioID2)^30^, (4) HA Tag, (5) 2X Stop codon, (6) a FRT site, and (7) a 1019 bp 3’ homology arm, beginning at the *Nphs2*-sgRNA cut site. The donor vector was designed to produce a final knock-in allele which would produce a fusion protein of podocin C-terminally linked to BioID2.

### Embryo microinjection

C57BL/6J zygotes were microinjected with 400nM Cas9 protein, 25 ng/μl sgRNA and 20 ng/μl supercoiled double-stranded donor plasmid (Mix1), or 200 nM Cas9 protein, 12.5 ng/μl sgRNA and 10 ng/μl donor plasmid (Mix2). Injected embryos were implanted in pseudopregnant B6D2F1 recipient females. Fourteen resulting pups (9 from Mix1 and 7 from Mix2) were screened by PCR for the presence of the knock-in allele. One female (Founder #2) and one male (Founder #6) were positive for the correct single-copy knock-in allele. Founders #2 and #6 were mated to wild-type C57BL/6J animals for transmission of the knock-in allele. Both founders transmitted the correct knock-in allele to offspring. Injections, genotyping of founders, and off-site targeting analysis was performed by UNC Chapel Hill Animal Models Core.

### Genotyping

A mouse tail or ear clip was taken and dissociated with Viagen DirectPCR Lysis Reagent (Mouse Tail) containing 10 µg/mL proteinase K incubated at 55°C overnight and denatured at 95 °C for 10 min. PCR was run with (T_annealing_ = 63.5°C), elongation for 40 sec, for 35 cycles. Primer sequences utilized for genotyping are Common Forward: 5’-CTTTTGTCCTCTCCCGGCAA-3’, podocin WT Reverse 5’-TGCATGTAGCCATCTTGTGACT-3’, *Nphs2*^*BioID2*^ Reverse 5’-CTGCCCTTGGTCTGTCTGTC-3’.

### Kidney cortex isolation and lysis

Mice were raised, housed, and handled in accordance with IACUC protocol number 19-183.0/22-136.0. 8–10-week-old mice were injected subcutaneously with 5 mg/kg biotin every day for one week. We surgically isolated the cortex of the kidney to enrich for the glomerular fraction, using a scalpel in cold sterile phosphate buffered saline (PBS). A single biological sample was composed of the isolated kidney cortex of three same sex mice for a total of six kidney cortexes per sample (2 kidneys per animal x 3 mice). The sample was then homogenized via a glass Dounce homogenizer. Samples were centrifuged at 4 °C, 5000 × g for 10 minutes. Supernatant was decanted and the tissue pellets snap frozen in liquid nitrogen and stored at -80 °C. Care was taken to utilize sterile Eppendorf tubes that had not been autoclaved as we identified a polyethylene glycol contaminant in a preliminary MS analysis that may arise from autoclaving plastic utilized. Samples were removed from the freezer and allowed to equilibrate on ice for 2 hours. Lysis buffer (8M urea, 50mM Tris-HCl Ph 7.4, 500mM NaCl, 2.5mM EDTA, 2.5mM EGTA, 1.5mM MgCl2, 1.5mM DTT, 0.25% NP-40, 1% SDS, 1x protease inhibitors (Roche cOmplete Mini EDTA-free, Sigma)) were then added to resuspend the pellet and allowed to nutate at 4°C for one hour. The protein homogenate was sonicated (3 pulses for 10 sec at 30% duty) and centrifuged (4°C, 12000 rpm for 10 min). The supernatant was then removed for subsequent quantification and analysis.

### Biotinylated protein capture

Kidney cortex-isolated protein lysates were serial diluted and run-in triplicate on a 96-well plate reader at 590nm wavelength (Synergy HT, BioTek). A standard curve of bovine serum albumin (BSA) was utilized as a control. Protein concentrations of each lysate were calculated, and 10mg of crude protein lysate was loaded with 100μl of streptavidin-coated magnetic beads (Dynabeads MyOne Streptavidin C1, Invitrogen). Samples containing beads and protein lysate were rotated end over end at 4°C overnight. Supernatant was removed using a magnetic strip, and the bead-captured fraction was washed once with lysis buffer. After the first wash with lysis buffer containing 1% SDS and 0.25 % NP-40 diminishing amounts of detergents were utilized in subsequent washes until there was no detergents remaining in the wash buffer (after 4 washes). Beads were then washed 3 times in ABC solution (50mM ammonium bicarbonate, 8.0 pH) and sent to UNC Chapel Hill Hooker Proteomics Core. All procedures were performed the same for our aged murine cohort with the exception that 5 mg of crude protein lysate was loaded onto beads.

After the last wash buffer step, 50 µl of 50mM ammonium bicarbonate (pH 8) containing 1 µg trypsin (Promega) was added to beads overnight at 37ºC with shaking. The next day, 500ng of trypsin was added then incubated for an additional 3h at 37ºC with shaking. Supernatants from pelleted beads were transferred, then beads were washed twice with 50ul LC/MS grade water. These rinses were combined with original supernatant, then acidified to 2% formic acid. Peptides were desalted with peptide desalting spin columns (Thermo Scientific) and dried via vacuum centrifugation. Peptide samples were stored at -80°C until further analysis.

### LC/MS/MS analysis

Each sample was analyzed by LC-MS/MS using an Easy nLC 1200 coupled to a QExactive HF (Thermo Scientific). Samples were injected onto an Easy Spray PepMap C18 column (75μm id × 25cm, 2μm particle size) (Thermo Scientific) and separated over a 120 min method. The gradient for separation consisted of a step gradient from 5 to 36 to 48% mobile phase B at a 250 nl/min flow rate, where mobile phase A was 0.1% formic acid in water and mobile phase B consisted of 0.1% formic acid in ACN. The QExactive HF was operated in data-dependent mode where the 15 most intense precursors were selected for subsequent HCD fragmentation. Resolution for the precursor scan (m/z 350–1700) was set to 60,000 with a target value of 3 × 10^6^ ions, 100ms inject time. MS/MS scans resolution was set to 15,000 with a target value of 1 × 10^5^ ions, 75ms inject time. The normalized collision energy was set to 27% for HCD, with an isolation window of 1.6m/z. Peptide match was set to preferred, and precursors with unknown charge or a charge state of 1 and ≥ 8 were excluded.

### Western blot

Biotinylated proteins attached to streptavidin coated bead were eluted off with excess biotin [200 mM] in 200 mM Tris HCl pH: 6.8, 40% glycerol, 8 % beta mercaptoethanol, 2 % SDS, 0.04 % bromophenol blue at 95°C for 30 min). Bead slurry mix was placed on a magnet and isolated biotinylated lysate was collected for subsequent analysis. Protein sample fractions were removed and loaded into a Novex WedgeWell gel (4-20% Tris-Glycine gradient gel, Invitrogen, XP04202), and run in 25mM Tris-HCl, 190mM Glycine, 0.1% SDS, pH 8.3. Proteins were transferred from gel to nitrocellulose membrane in 25mM Tris-HCl, 190mM Glycine, and 20% methanol. The membrane was blocked in 3% bovine serum albumin in 1x TBST (Tris base Saline Solution (25mM Tris-HCl pH 7.5, 150mM NaCl) with 0.1% Tween-20, for 1 hour at room temperature. Primary antibodies (Rabbit anti-HA tag (Cell Signaling, 3724S [1:500], mouse anti-NPHS2 (Proteintech, 20384-1-AP) [1:500], Rabbit anti-podocin (Invitrogen, PA5-79757) [1:500], Streptavidin-HRP (Cell Signaling, 3999S) [1:1000] were applied in 3% BSA+ TBST. Membrane plus primary antibodies were allowed to incubate overnight at 4°C with gentle nutation. Following 3×10 min washes with TBST, membranes were probed with secondary antibodies conjugated to horseradish peroxidase (HRP) (donkey and rabbit -HRP [1:500] or Goat anti mouse-HRP [1:500] and incubated for one hour at room temperature Membranes were developed with enhanced chemiluminescence substrate (ECL) and visualized on an iBright FL1000 (Invitrogen).

### Proteomics data analysis

Raw data were processed using the MaxQuant software suite (version 1.6.12.0) for identification and label-free quantitation.^62^ Data were searched against an Uniprot Reviewed Mouse database (downloaded January 2021, containing 17,051 sequences) using the integrated Andromeda search engine. A maximum of two missed tryptic cleavages was allowed. The variable modification specified was oxidation of methionine, N-terminal acetylation, and phosphorylation of Ser/Thr/Tyr. Label-free quantitation (LFQ) was enabled. Results were filtered to 1% FDR at the unique peptide level and grouped into proteins within MaxQuant. Match between runs was enabled. Data were filtered in Perseus, then imported into Argonaut for normalization, imputation and statistical analysis.^63^ We combined three separate MS analyses and averaged the Log_2_ fold changes and respective *p*-values of the top 17 proteins identified across all three MS profiles. QIAGEN Ingenuity Pathway Analysis (IPA) was utilized to identify putative interactomes.

### Immunofluorescence

Kidneys were harvested in cold filter sterilized PBS and fixed at 4°C for 1 hr in 4% paraformaldehyde (PFA) in PBS. Samples were washed twice in 1x PBS, placed in 30% sucrose overnight, and subsequently embedded in OCT. Kidneys were cut in 12μm sections on a Leica Cryostat CM 1850. Tissue sections were blocked in 3% donkey serum, 1% bovine serum albumin (BSA), 1x PBS, 0.1 % TritonX-100 for 45 min at room temperature. Slides were incubated in primary antibodies (see antibodies utilized below, Supplemental Table I) diluted in blocking buffer for 1-2 hr. Slides were then rinsed 3×5 min in 1x PBST, after which they were incubated with Alexa Fluor labeled secondary antibodies (Invitrogen) diluted (1:1000) in blocking buffer for 1 hr at room temp. The slides were then rinsed 2 × 5 min with 1x PBST, 1×3–5 min with 1x PBS + 1 ng/mL DAPI and mounted in ProLong Gold antifade reagent (Invitrogen).

For double labeling tissue with two antibodies both raised in rabbit we utilized a Zenon double labeling kit (Z25302, Invitrogen) with rabbit anti-Fnbp1l, rabbit anti-Ildr2, rabbit anti-podocin, and rabbit anti-HA antibodies. The weaker of the two primary rabbit antibodies, typically Ildr2 and Fnbp1l, was diluted in block solution then incubated for 90 min at room temperature. Slides were washed 3-6x with 1xPBS +0.25% Triton-X 100 and subsequently incubated for 1hr at room temperature with the appropriate Alexa Fluor-488 labeled secondary antibody diluted 1:1000 in blocking buffer. The second primary rabbit antibody was mixed at a 1:2.5 ratio with the Zenon fluorophore-647. Note the two fluorophores utilized are on opposite ends of the florescence spectrum. The slides were then fixed in 4% PFA for 5 min at room temperature. And all subsequent steps were carried out per normal immunofluorescence procedures as above. Images were acquired utilizing a Zeiss 880 confocal microscope equipped with Airyscan super-resolution and spectral imaging on Zen Microscopy Suite version 2.3 Sp1 that is part of the UNC Hooker Imaging Core. Z-stack images were acquired in 1μm steps for Ildr2 and Fnbp1l.

### Immunogold electron microscopy

Mouse kidneys were fixed for 1 hour at room temperature in 4% PFA in 0.15M sodium phosphate buffer pH 7.4 (PB) or 2% PFA + 0.5% glutaraldehyde in 0.15M PB and immediately processed for LR White resin embedding. Samples were washed in 0.15M PB, 3×10 minutes and dehydrated using a Pelco BioWave Pro Microwave (Ted Pella, Inc.) as follows: at 40°C (750 watts): 30% ethanol in water (ETOH) -1 min, 50% ETOH in water-1 min, 75% ETOH in water-1 min, and at 40°C (450 watts): 100% ETOH-1 min, 100% ETOH-1 min, 100% ETOH-1 min. Microwave infiltration and embedment were carried out using the following schedule: 1 part 100% ETOH:2 part LR White resin-10 min at 40°C (350 watts), 2 exchanges of 100% LR White resin for 10 min at 50°C (350 watts). Samples were transferred to 00 gelatin capsules filled with fresh LR White resin and polymerized overnight at 55°C; the temperature was adjusted to 60°C for 6 hours and allowed to complete polymerization at room temperature for 72 hrs. Blocks were trimmed to the tissue and 1.0µm sections were cut using Leica Ultracut UCT (Leica Microsystems, Inc.) and a Diatome diamond knife (Electron Microscopy Sciences). Sections were mounted on glass slides, stained with 1% toluidine blue O in 1% sodium borate, and regions with glomeruli were selected and trimmed.^64^ Ultrathin sections (90 nm) were cut and mounted on formvar / carbon-coated 200 mesh nickel grids. Before immunostaining, the sections were hydrated by floating the grids section-side down on drops of deionized water. Sections were blocked with Aurion Goat Blocking Solution for 15 min and transferred to 15µl drops of the primary antisera diluted at [1: 500] for anti-HA and [1: 250] for anti-Podocin in 0.05M TBS+0.2% BSA-Ac, pH 7.6 (Rabbit anti-HA, Cell Signaling, 37245; rabbit anti-Podocin, Invitrogen, PA5-79757). Sections were incubated overnight at 4°C followed by 4×10 min washes in TBS/BSA-Ac to remove unbound antibody. The grids were incubated in a 12nm Colloidal gold-AffiniPure goat anti-rabbit IgG (H+L) secondary antibody (Jackson Immuno, Lot #148041) diluted at 1:50 in TBS/BSA-Ac for 2 hours at room temperature.^65^ After 3 washes in TBS/BSA-Ac and 3 washes in 0.15M PB, grids were post-fixed for 10 min in 1% glutaraldehyde in 0.15M PB followed by 3 washes in deionized water. The grids were stained with 4% aqueous uranyl acetate for 5 min for additional contrast. Samples were observed using a JEOL JEM-1230 transmission electron microscope operating at 80kV (JEOL USA INC.), and images were taken using a Gatan Orius SC1000 CCD camera with Gatan Microscopy Suite version 3.10.1002.0 software (Gatan, Inc.). Sample prep and imaging was performed at the UNC Microscopy Services Laboratory, Department of Pathology and Laboratory Medicine.

### In situ hybridization

Wildtype C57/Bl6J kidneys at postnatal day 2 (P2) were dissected in cold molecular grade PBS (RNAse/DNAse free) and fixed in 4% PFA for 30 min at room temperature on a rocking platform. Kidneys were washed in molecular grade PBS and placed in 30% sucrose. Kidneys were embedded in OCT and 20μm sections were cut on a cryostat (Leica CM1850). Tissue sections were fixed in 4% PFA and washed with 1x PBS. The tissue was permeabilized with proteinase K at 10μg/mL for 15 min and subsequently washed with 1x PBS. Tissue was then washed in an acetylation solution (0.1 % HCl, 0.375 % Acetic Anhydride, and 0.75 % Triethanolamine in H_2_0) for 10 min with stirring. Tissue sections were then subsequently washed 3×3min with PBS, rinsed 1×5min in 0.85 % NaCl, 1×5min in 70 % ethanol, and 1×5min in 95 % ethanol prior to riboprobe / hybridization application. Antisense Digoxygenin (DIG) labeled riboprobes were hybridized to the tissue in a solution containing 50% deionized formamide, 2 sodium citrate pH 4.5, 1% SDS, 50μg/mL heparin, 50μg/mL yeast tRNA overnight in a humidity chamber at 68°C. The following day, specimens were treated with successive washes of sodium citrate buffer as described previously,^66^ and blocked in 10% heat inactivated sheep serum (HISS), with 2% Roche blocking reagent (BR) in malic acid buffered solution with 0.1% tween-20 (MABT). Anti-DIG-Alkaline Phosphatase (AP) antibody was applied in 1% HISS, 2% BR in MABT overnight at 4°C. The following day the slides were washed in MABT and 100mM NaCl, 100 mM Tris-HCl pH 9.5, 50mM MgCl_2_, and 0.1 % Tween-20 (NTMT) with 2mM Levamisole to inhibit endogenous alkaline phosphatase activity. Digoxigenin-UTP (Roche 53119620) labeled riboprobes were amplified from cDNA libraries collected from mouse kidney tissue with the following primers:

Podocin forward primer: TGACGTTCCCTTTTTCCATC, Podocin reverse primer with T7 underlined: CAGTGAATTG<u>TAATACGACTCACTATAGGG</u>CTGTGGACAGCGACTGAAGA. Ildr2 forward: GGAGAATCCTTGGGC and Ildr2 reverse: CAGTGAATTG<u>TAATACGACTCACTATAGGGG</u>TACCCGGCCTTGGC were previously published^66^. Fnbp1l forward primer: GCTGAATGACAATTGTGTGAAC and Fnbp1l reverse: CAGTGAATTG<u>TAATACGACTCACTATAGGG</u>CTGTGCAAGTCCAAGTGTCTTC.

### Human kidney tissue

Human kidney tissue samples were de-identified (young) or directly donated (91-year-old) and did not necessitate IRB approval. Young, normal kidney tissue was obtained from the UNC Tissue Procurement Facility as a frozen block. A small piece of tissue was removed with a razorblade, fixed for 10 minutes in 4% paraformaldehyde and subsequent processing for cryosectioning and immunofluorescence carried out as for the mouse (described above). For the old kidney tissue, a fresh sample was excised from the kidney cortex and fixed for 10 minutes in 4% paraformaldehyde prior to processing similar to the young tissue for immunofluorescence.

## Acknowledgements

We would like to acknowledge the many people and core services at the University of North Carolina at Chapel Hill who aided in this investigation. We specifically would like to thank Wendy Salmon, Director of the Department of Cell Biology and Physiology’s Hooker Imaging Core, Laura Herring, Ph.D., Director of the School of Medicine’s Michael Hooker Proteomics Center, Victoria Madden and Kristen White, providers of electron microscopy services in the Department of Pathology and Laboratory Medicine’s Microscopy Services Laboratory, and Paul Risteff for training in TEM and aid with image acquisition and sample preparation. We would further like to acknowledge the members of the O’Brien lab for critical feedback on these studies.

This research is based in part on work conducted at the Microscopy Services Laboratory (TEM), the UNC Proteomics Core Facility (MS), and the UNC Hooker Imaging Core Facility (confocal microscopy), which are supported in part by P30 CA016086 Cancer Center Core Support Grant to the UNC Lineberger Comprehensive Cancer Center. This research was supported in part by a Vanderbilt O’Brien Kidney Center Pilot and Feasibility Award (P30-DK114809), a UNC Junior Faculty Development Award, and Start-Up funds from UNC to LLO as well as F32AR073649 to GFG.

## Author contributions

G.F.G. helped design the study, performed the experiments, and wrote the manuscript. Z.H.I., S.L.C., and A.N.S. helped perform the experiments. L.L.O. conceptualized the podocin-BioID model, designed and oversaw the study, and edited the manuscript.

