## Supplemental Table and Figures for "Mapping of the podocin proximity-dependent proteome reveals novel components of the kidney podocyte foot process"

| Antidody | Dilution [IF] | Dilution [WB] | Company |
|---|---|---|---|
| Goat anti - Collagen IV | [1:500] | | Sigma-Aldrich, AB769, Lot 2930544 |
| Mouse (IgG1) anti - Myozap | supernatant (straight) | | Progen, clone 517.67, 651169 |
| Rabbit anti - F11r (Jam1) | [1:50] | | ABclonal, A1241, Lot 0005180101 |
| Rabbit anti - Fnbp1l (Toca-1) | [1:100] | | Bethyl Laboratories, A303-469A, Lot 1 |
| Rabbit anti - HA | [1:500] | [1:1000] | Cell Signaling, clone C29F4, 3724S |
| Rabbit anti - Ildr2 | [1:250] | | BiCell Scientific Inc, 00303 |
| Rabbit anti - Ildr2 | [1:50] | | Lifespan BioSciences LS-C205504 |
| Rabbit anti - Ildr2 | [1:50] | | Invitrogen, PA5-46481, Lot WE327570 |
| Rabbit anti - Pard3 | [1:250] | | Sigma-Aldrich, 07-330, Lot 2615671 |
| Rabbit anti - podocin | [1:500] | [1:1000] | Invitrogen, PA5-79757, Lot WL3458408 |
| Rabbit anti - WT1 | [1:500] | | Abcam, ab89901, LotGR3365362-7 |
| Lotus Lectin | [1:1000] | | Vector Laboratories, FL-1321, Lot ZC0914 |
| Streptavidin - APC | [1:1000] | | BD Pharmingen, 554067 Lot 7040924 |
| Streptavidin - HRP | | [1:5000] | Cell Signaling, 3999S, Lot 8 |

**Supplemental Table I. Antibodies** List of antibodies with respective company identifiers and concentrations for immunoflorecence (IF) and western blot (WB) analyses utilized in this investigation.

| Gene Symbol | Majority Protein IDs | Protein Names | Razor + Unique Peptides | Log2 Fold Change | *p* - Value |
|---|---|---|---|---|---|
| *Nphs2* | Q91X05 | Podocin | 12 | 10.63 | 0.00003 |
| *Tjp1* | P39447 | Tight junction protein ZO-1 | 85 | 7.40 | 0.001 |
| *Magi2* | Q9WVQ1 | Membrane-associated guanylate kinase, WW and PDZ domain-containing protein 2 | 33 | 7.37 | 0.001 |
| *Kirrel* | Q80W68 | Kin of IRRE-like protein 1 | 20 | 7.06 | 0.001 |
| *Pard3b* | Q9CSB4 | Partitioning defective 3 homolog B | 32 | 5.34 | 0.00002 |
| *Ddn* | Q80TS7 | Dendrin | 12 | 5.33 | 0.00003 |
| *Ildr2* | B5TVM2 | Immunoglobulin-like domain-containing receptor 2 | 11 | 4.51 | 0.009 |
| *Cttn* | Q60598 | Src substrate cortactin | 15 | 4.01 | 0.0003 |
| *Nebl* | Q9DC07 | LIM zinc-binding domain-containing Nebulette | 11 | 3.65 | 0.003 |
| *Tjp2* | Q9Z0U1 | Tight junction protein ZO-2 | 31 | 3.63 | 0.021 |
| *S100g* | P97816 | Protein S100-G | 4 | 3.06 | 0.034 |
| *Fnbp1l* | Q8K012 | Formin-binding protein 1-like | 7 | 2.92 | 0.010 |
| *Capza2* | P47754 | F-actin-capping protein subunit alpha-2 | 8 | 2.82 | 0.014 |
| *Atp6v1e1* | P50518 | V-type proton ATPase subunit E 1 | 9 | 2.30 | 0.039 |
| *Canx* | P35564 | Calnexin | 9 | 2.26 | 0.042 |
| *Myzap* | Q3UIJ9 | Myocardial zonula adherens protein | 7 | 2.09 | 0.0001 |
| *Kpnb1* | P70168 | Importin subunit beta-1 | 2 | 1.96 | 0.041 |
| *Dars* | Q922B2 | Aspartate--tRNA ligase, cytoplasmic | 7 | 1.88 | 0.029 |
| *Cndp2* | Q9D1A2 | Cytosolic non-specific dipeptidase | 6 | 1.83 | 0.041 |
| *Pls3* | Q99K51 | Plastin-3 | 5 | 1.76 | 0.029 |
| *Sult1d1* | Q3UZZ6 | Sulfotransferase 1 family member D1 | 5 | 1.73 | 0.038 |
| *Uqcr10* | Q8R1I1 | Cytochrome b-c1 complex subunit 9 | 3 | 1.67 | 0.048 |
| *Uba52* | P62984; P62983; P0CG49; P0CG50 | Ubiquitin-60S ribosomal protein L40 | 5 | 1.60 | 0.041 |
| *Sec23a* | Q01405 | Protein transport protein Sec23A | 7 | 1.59 | 0.030 |
| *Cisd2* | Q9CQB5 | CDGSH iron-sulfur domain-containing protein 2 | 2 | 1.46 | 0.026 |
| *Naca* | Q60817;P70670 | Nascent polypeptide-associated complex subunit alpha, muscle-specific form | 2 | 1.46 | 0.049 |
| *Ca15* | Q99N23 | Carbonic anhydrase 15 | 3 | 1.44 | 0.050 |
| *Qars* | Q8BML9 | Glutaminyl-TRNA Synthetase 1 | 2 | 1.40 | 0.039 |
| *Rpl35* | Q6ZWV7 | 60S ribosomal protein L35 | 4 | 1.37 | 0.038 |
| *Agk* | Q9ESW4 | Acylglycerol kinase, mitochondrial | 5 | 1.37 | 0.039 |
| *Scp2* | P32020 | Non-specific lipid-transfer protein | 13 | 1.36 | 0.026 |
| *Aimp1* | P31230 | Endothelial monocyte-activating polypeptide 2 | 2 | 1.26 | 0.046 |
| *Dhrs7b* | Q99J47 | Dehydrogenase/reductase SDR family member 7B | 5 | 1.24 | 0.024 |
| *Fabp4* | P04117 | Fatty acid-binding protein, adipocyte | 2 | 1.17 | 0.012 |
| *Mpv17* | P19258 | Protein Mpv17 | 2 | 1.15 | 0.007 |
| *Sf3b1* | Q99NB9 | Splicing factor 3B subunit 1 | 4 | 1.10 | 0.039 |
| *Dkc1* | Q9ESX5 | H/ACA ribonucleoprotein complex subunit 4 | 4 | 1.06 | 0.003 |
| *Por* | P37040 | NADPH--cytochrome P450 reductase | 11 | 1.05 | 0.031 |
| *Synpo* | Q8CC35 | Synaptopodin | 6 | 1.02 | 0.054 |
| *Acox1* | Q9R0H0 | Peroxisomal acyl-coenzyme A oxidase 1 | 24 | 1.01 | 0.015 |
| *Cisd1* | Q91WS0 | CDGSH iron-sulfur domain-containing protein 1 | 6 | 0.99 | 0.036 |

**Supplemental Table II. Proteomic profiling of cohort 1, 8–10-week-old *Nphs2*$^{BioID2/+}$ male mice, identifies 41 enriched podocyte foot process proteins.** MS analysis of biotin administered male *Nphs2*$^{BioID2/+}$ mice identifies podocin (Nphs2), Tjp1, Magi2, Kirrel, and Pard3b as the top 5 proteins, respectively. Table lists 41 significant proteins ($p \leq 0.05$) at a $\log_2$ cut off 1.0 with their respective *p*-values in the far-right column. Podocin was identified as the top hit and is documented to oligomerize with itself supporting the efficacy of the model. All proteins were identified with at least 2 razor unique peptide sequences *i.e*., two peptide sequences aligned only to that protein.

| Gene Symbol | Majority Protein IDs | Protein Names | Razor + Unique Peptides | $Log_2$ Fold Change | *p* - Value |
|---|---|---|---|---|---|
| *Kirrel* | Q80W68 | Kin of IRRE-like protein 1+D3:D62 | 15 | 10.19 | 0.001 |
| *Tjp1* | P39447 | Tight junction protein ZO-1 | 85 | 9.88 | 0.003 |
| *Pard3b* | Q9CSB4 | Partitioning defective 3 homolog B | 26 | 9.62 | 0.016 |
| *Ddn* | Q80TS7 | Dendrin | 11 | 8.99 | 0.005 |
| *Magi2* | Q9WVQ1 | Membrane-associated guanylate kinase, WW and PDZ domain-containing protein 2 | 24 | 8.61 | 0.011 |
| *Nebl* | Q9DC07 | LIM zinc-binding domain-containing Nebulette | 7 | 7.75 | 0.021 |
| *Ildr2* | B5TVM2 | Immunoglobulin-like domain-containing receptor 2 | 9 | 7.42 | 0.010 |
| *Cttn* | Q60598 | Src substrate cortactin | 14 | 7.08 | 0.020 |
| *Nphs2* | Q91X05 | Podocin | 10 | 7.05 | 0.023 |
| *Fnbp1l* | Q8K012 | Formin-binding protein 1-like | 5 | 6.65 | 0.008 |
| *Magi1* | Q6RHR9 | Membrane-associated guanylate kinase, WW and PDZ domain-containing protein 1 | 6 | 5.89 | 0.002 |
| *Tjp2* | Q9Z0U1 | Tight junction protein ZO-2 | 30 | 5.78 | 0.006 |
| *Myzap* | Q3UIJ9 | Myocardial zonula adherens protein | 4 | 4.80 | 0.018 |
| *Synpo* | Q8CC35 | Synaptopodin | 4 | 4.79 | 0.043 |
| *Khsrp* | Q3U0V1 | Far upstream element-binding protein 2 | 5 | 4.69 | 0.014 |
| *Mapre1* | Q61166 | Microtubule-associated protein RP/EB family member 1 | 4 | 4.56 | 0.019 |
| *Yes1* | Q04736 | Tyrosine-protein kinase Yes | 8 | 4.34 | 0.015 |
| *Tmem65* | Q4VAE3 | Transmembrane protein 65 | 4 | 4.25 | 0.030 |
| *Cd2ap* | Q9JLQ0 | CD2-associated protein | 5 | 4.15 | 0.019 |
| *Ap1g1* | P22892 | AP-1 complex subunit gamma-1 | 6 | 3.85 | 0.018 |
| *Ewsr1* | Q61545 | RNA-binding protein EWS | 2 | 3.83 | 0.038 |
| *Oxsm* | Q9D404 | 3-oxoacyl-[acyl-carrier-protein] synthase, mitochondrial | 4 | 3.77 | 0.033 |
| *Erlin2* | Q8BFZ9 | Erlin-2 | 6 | 3.44 | 0.008 |
| *Epb41l5* | Q8BGS1 | Band 4.1-like protein 5 | 9 | 3.13 | 0.003 |
| *Hsd3b4* | Q61767 | 3 beta-hydroxysteroid dehydrogenase type 4 | 9 | 2.99 | 0.028 |
| *Dhtkd1* | A2ATU0 | Probable 2-oxoglutarate dehydrogenase E1 component DHKTD1, mitochondrial | 7 | 2.93 | 0.017 |
| *Taco1* | Q8K0Z7 | Translational activator of cytochrome c oxidase 1 | 2 | 2.64 | 0.045 |
| *Aif1l* | Q9EQX4 | Allograft inflammatory factor 1-like | 3 | 2.31 | 0.001 |
| *Gstt2* | Q61133 | Glutathione S-transferase theta-2 | 4 | 2.24 | 0.002 |
| *Sorbs1* | Q62417 | Sorbin and SH3 domain-containing protein 1 | 18 | 2.21 | 0.036 |
| *Tns2* | Q8CGB6 | Tensin-2 | 12 | 2.20 | 0.007 |
| *Farp1* | F8VPU2 | FERM, RhoGEF and pleckstrin domain-containing protein 1 | 7 | 2.13 | 0.040 |
| *Tjp3* | Q9QXY1 | Tight junction protein ZO-3 | 7 | 2.12 | 0.043 |
| *Slc25a42* | Q8R0Y8 | Mitochondrial coenzyme A transporter SLC25A42 | 9 | 1.98 | 0.021 |
| *Mybbp1a* | Q7TPV4 | Myb-binding protein 1A | 13 | 1.92 | 0.037 |
| *Acnat1* | A2AKK5 | Acyl-coenzyme A amino acid N-acyltransferase 1 | 4 | 1.81 | 0.046 |
| *Epb41l1* | Q9Z2H5 | Band 4.1-like protein 1 | 8 | 1.79 | 0.023 |
| *Col1a1* | P11087 | Collagen alpha-1(I) chain | 8 | 1.73 | 0.002 |
| *Col4a2* | P08122 | Collagen alpha-2(IV) chain;Canstatin | 5 | 1.69 | 0.006 |
| *Nomo1* | Q6GQT9 | Nodal modulator 1 | 17 | 1.68 | 0.014 |
| *Col1a2* | Q01149 | Collagen alpha-2(I) chain | 10 | 1.65 | 0.001 |
| *Dao* | P18894 | D-amino-acid oxidase | 11 | 1.63 | 0.002 |
| *Arpc4* | P59999 | Actin-related protein 2/3 complex subunit 4 | 2 | 1.62 | 0.007 |
| *Uqcrh* | P99028 | Cytochrome b-c1 complex subunit 6, mitochondrial | 2 | 1.61 | 0.019 |
| *Atp5l* | Q9CPQ8 | ATP synthase subunit g, mitochondrial | 4 | 1.59 | 0.002 |
| *Patj* | Q63ZW7 | Pals1-associated tight junction protein / InaD-like protein | 6 | 1.53 | 0.003 |
| *Dhrs1* | Q99L04 | Dehydrogenase/reductase SDR family member 1 | 8 | 1.53 | 0.0003 |
| *Apmap* | Q9D7N9 | Adipocyte plasma membrane-associated protein | 14 | 1.52 | 0.003 |
| *Lad1* | P57016 | Ladinin-1 | 14 | 1.49 | 0.012 |
| *Rtn4* | Q99P72 | Reticulon-4 | 8 | 1.49 | 0.012 |
| *F11r* | O88792 | Junctional adhesion molecule A | 2 | 0.70 | 0.050 |

**Supplemental Table III. Proteomic profiling of cohort 2, 8–10-week-old *Nphs2*$^{BioID2/+}$ male mice, identifies 50 enriched podocyte foot process proteins.** MS analysis of *Nphs2*$^{BioID2/+}$ biotin administered mice identifies Kirrel, Tjp1, Pard3b, Ddn, and Magi2 as the top 5 proteins detected respectively, with podocin detected in the top 10. Table lists 50 significant proteins ($p \leq 0.05$) at a $\log_2$ cut off $\geq$ 1.49 with respective *p*-values in the far-right column. One additional immunoglobulin domain containing protein, Junction adhesion molecule 1 (Jam1 / F11r) was also identified within this MS analysis with a $Log_2$ fold change of 0.7, $p \leq 0.05$. All proteins were detected with at least two razor unique peptide sequences to the specific protein identified.

| Gene Symbol | Majority Protein IDs | Protein Names | Razor + Unique Peptides | $Log_2$ Fold Change | *p* - Value |
|---|---|---|---|---|---|
| *Kirrel* | Q80W68 | Kin of IRRE-like protein 1 | 18 | 5.99 | 0.027 |
| *Cttn* | Q60598 | Src substrate cortactin | 14 | 5.70 | 0.034 |
| *Tjp1* | P39447 | Tight junction protein ZO-1 | 90 | 5.62 | 0.050 |
| *Pard3b* | Q9CSB4 | Partitioning defective 3 homolog B | 38 | 4.21 | 0.050 |
| *Nebl* | Q9DC07 | LIM zinc-binding domain-containing Nebulette | 7 | 3.41 | 0.051 |
| *Fnbp1l* | Q8K012 | Formin-binding protein 1-like | 7 | 3.22 | 0.025 |
| *Magi2* | Q9WVQ1 | Membrane-associated guanylate kinase, WW and PDZ domain-containing protein 2 | 30 | 2.89 | 0.044 |
| *Synpo* | Q8CC35 | Synaptopodin | 10 | 2.88 | 0.044 |
| *Ildr2* | B5TVM2 | Immunoglobulin-like domain-containing receptor 2 | 11 | 2.76 | 0.003 |
| *Tjp2* | Q9Z0U1 | Tight junction protein ZO-2 | 21 | 2.71 | 0.037 |
| *Col1a2* | Q01149 | Collagen alpha-2(I) chain | 3 | 2.68 | 0.046 |
| *Nphs2* | Q91X05 | Podocin | 13 | 2.42 | 0.035 |
| *Col1a1* | P11087 | Collagen alpha-1(I) chain | 5 | 2.37 | 0.011 |
| *Ddn* | Q80TS7 | Dendrin | 13 | 2.35 | 0.021 |
| *Pkp4* | Q68FH0 | Plakophilin-4 | 13 | 2.27 | 0.033 |
| *Cd2ap* | Q9JLQ0 | CD2-associated protein | 3 | 2.20 | 0.034 |
| *Rdx; Msn* | P26043; P26041 | Radixin; Moesin | 2 | 1.74 | 0.015 |
| *Afdn* | Q9QZQ1 | Afadin | 2 | 1.70 | 0.030 |
| *Hspe1* | Q64433 | 10 kDa heat shock protein, mitochondrial | 4 | 1.66 | 0.036 |
| *Aif1l* | Q9EQX4 | Allograft inflammatory factor 1-like | 5 | 1.42 | 0.027 |
| *Hist1h4a* | P62806 | Histone H4 | 9 | 1.36 | 0.014 |
| *Tns2* | Q8CGB6 | Tensin-2 | 3 | 1.30 | 0.038 |
| *Khk* | P97328 | Ketohexokinase | 2 | 1.25 | 0.027 |
| *Clic1* | Q9Z1Q5 | Chloride intracellular channel protein 1 | 2 | 1.21 | 0.027 |
| *Hist2h2bb* | Q8CGP2;Q8CGP1 | Histone H2B type 1 | 6 | 1.04 | 0.049 |
| *Acadl* | P51174 | Long-chain specific acyl-CoA dehydrogenase, mitochondrial | 7 | 1.01 | 0.025 |

**Supplemental Table IV. Proteomic profiling of cohort 3, 8–10-week-old *Nphs2*$^{BioID2/+}$ female mice, identifies 28 enriched podocyte foot process proteins.** MS analysis of biotin administered female *Nphs2*$^{BioID2/+}$ mice identifies Kirrel, Cttn, Tjp1, Pard3b, and Nebl as the top 5 proteins detected**.** Table lists 28 significant proteins ($p \leq 0.05$) at a $\log_2$ cut off 1.0 with respective *p*-values in the far-right column. All proteins were detected with at least two razor unique peptide sequences to the specific protein identified.

| Gene Symbol | Protein Names | *p*-Value | Log 2 Fold Change |
|---|---|---|---|
| *Kirrel* | Kin of IRRE-like protein 1 | 0.0006 | 7.75 |
| *Tjp1* | Tight junction protein ZO-1 | 0.002 | 7.63 |
| *Nphs2* | Podocin | 0.019 | 6.70 |
| *Pard3b* | Partitioning defective 3 homolog B | 0.008 | 6.39 |
| *Magi2* | Membrane Associated Guanylate Kinase, WW And PDZ Domain Containing 2 | 0.018 | 6.29 |
| *Cttn* | Src substrate cortactin | 0.001 | 5.60 |
| *Ddn* | Dendrin | 0.003 | 5.56 |
| *Nebl* | LIM zinc-binding domain-containing Nebulette | 0.010 | 4.93 |
| *Ildr2* | Immunoglobulin-like domain-containing receptor 2 | 0.007 | 4.90 |
| *Fnbp1l* | Formin-binding protein 1-like | 0.014 | 4.27 |
| *Synpo* | Synaptopodin | 0.047 | 2.90 |
| *Tjp2* | Tight junction protein ZO-2 | 0.014 | 4.71 |
| *Myzap* | Myocardial zonula adherens protein | 0.009 | 3.45 |
| *Cd2ap* | CD2-associated protein | 0.020 | 3.18 |
| *Col1a1* | Collagen alpha-1(I) chain | 0.007 | 2.05 |
| *Aif1l* | Allograft inflammatory factor 1-like | 0.014 | 1.86 |
| *Tns2* | Tensin-2 | 0.022 | 1.75 |
| *Magi1* | Membrane Associated Guanylate Kinase, WW And PDZ Domain Containing 2 | 0.002 | 5.89 |
| *Khsrp* | Far upstream element-binding protein 2 | 0.014 | 4.69 |
| *Mapre1* | Microtubule-associated protein RP/EB family member 1 | 0.019 | 4.56 |
| *Yes1* | Tyrosine-protein kinase Yes | 0.015 | 4.34 |
| *Tmem65* | Transmembrane protein 65 | 0.030 | 4.25 |
| *Ap1g1* | AP-1 complex subunit gamma-1 | 0.018 | 3.85 |
| *Erlin2* | Erlin-2 | 0.008 | 3.44 |
| *Epb41l5* | Band 4.1-like protein 5 | 0.003 | 3.13 |
| *S100g* | Protein S100-G | 0.034 | 3.06 |
| *Capza2* | F-actin-capping protein subunit alpha-2 | 0.014 | 2.82 |
| *Atp6v1e1* | V-type proton ATPase subunit E 1 | 0.039 | 2.30 |
| *Pkp4* | Plakophilin-4 | 0.033 | 2.27 |
| *Canx* | Calnexin | 0.042 | 2.26 |
| *Sorbs1* | Sorbin and SH3 domain-containing protein 1 | 0.036 | 2.21 |
| *Farp1* | FERM, RhoGEF and pleckstrin domain-containing protein 1 | 0.040 | 2.13 |
| *Tjp3* | Tight junction protein ZO-3 | 0.043 | 2.12 |
| *Mybbp1a* | Myb-binding protein 1A | 0.037 | 1.92 |
| *Cndp2* | Cytosolic non-specific dipeptidase | 0.041 | 1.83 |
| *Epb41l1* | Band 4.1-like protein 1 | 0.023 | 1.79 |
| *Pls3* | Plastin-3 | 0.029 | 1.76 |
| *Rdx* | Radixin | 0.015 | 1.74 |
| *Msn* | Moesin | 0.015 | 1.74 |
| *Afdn* | Afadin | 0.021 | 1.70 |
| *Col4a2* | Collagen alpha-2(IV) chain;Canstatin | 0.006 | 1.69 |
| *Nomo1* | Nodal modulator 1 | 0.014 | 1.68 |
| *Col1a2* | Collagen alpha-2(I) chain | 0.0009 | 1.65 |
| *Arpc4* | Actin-related protein 2/3 complex subunit 4 | 0.007 | 1.62 |
| *Sec23a* | Protein transport protein Sec23A | 0.030 | 1.59 |
| *Patj* | Pals 1-associated tight junction protein / InaD-like protein | 0.003 | 1.53 |
| *Lad1* | Ladinin-1 | 0.012 | 1.49 |
| *Rtn4* | Reticulon-4 | 0.012 | 1.49 |
| *Cisd2* | CDGSH iron-sulfur domain-containing protein 2 | 0.026 | 1.46 |
| *Naca* | Nascent polypeptide-associated complex subunit alpha | 0.049 | 1.46 |
| *Scp2* | Non-specific lipid-transfer protein | 0.026 | 1.36 |
| *Aimp1* | Endothelial monocyte-activating polypeptide 2 | 0.046 | 1.26 |
| *Dhrs7b* | Dehydrogenase/reductase SDR family member 7B | 0.024 | 1.24 |
| *Clic1* | Chloride intracellular channel protein 1 | 0.027 | 1.21 |

**Supplemental Table V. Compiled proteomic profile of three cohorts of 8–10-week-old male and female *Nphs2*$^{BioID2/+}$ mice.** Table lists 54 significant proteins ($p \leq 0.05$) at a $\log_2$ cut off $\geq$ 1.20, with respective *p*–value listed in the far-right column. Table compiles the proteomic profiles across all three separate analyses and averages their respective $\log_2$ fold change and *p*-values. This table was subsequently utilized for *Qiagen Ingenuity Pathway Analysis* (IPA) and input into the *Database for Annotation, Visualization, and Integrated Discovery* (DAVID) for gene ontogeny characterization. All proteins were detected with at least 2 razor unique peptides specific to the protein denoted.

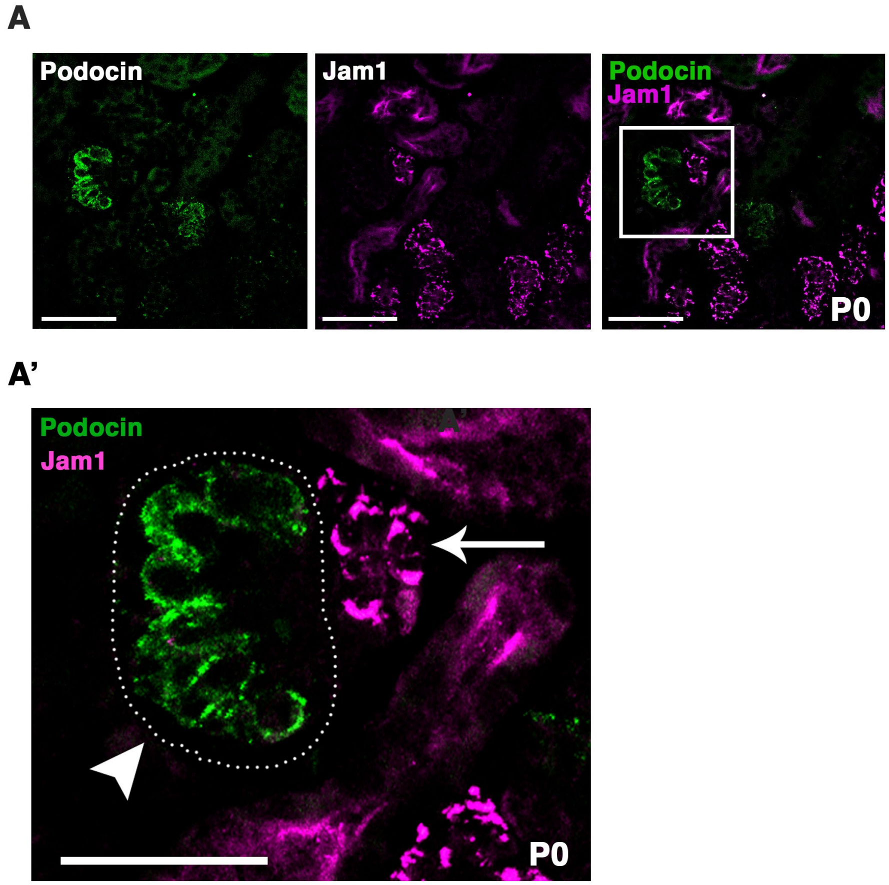


**Supplemental Figure 1. Expression of immunoglobulin superfamily member Jam1/F11r in tubule cells neighboring podocytes. (A)**. Jam1 was identified with a 0.7 $\log_2$ fold change in a single mass spec analysis. P0 kidney sections were immunostained for Jam1 (magenta) and podocin (green). Jam1 is localized to tubule cells directly adjacent to podocytes. **(A')** Highlighted white boxed region from **(A)** is enlarged to show podocin localization restricted to the glomerulus, outlined in a dotted while circle, and Jam1 localization in neighboring tubule cells, denoted by white arrow. All scale bars: 50 $\mu$m.

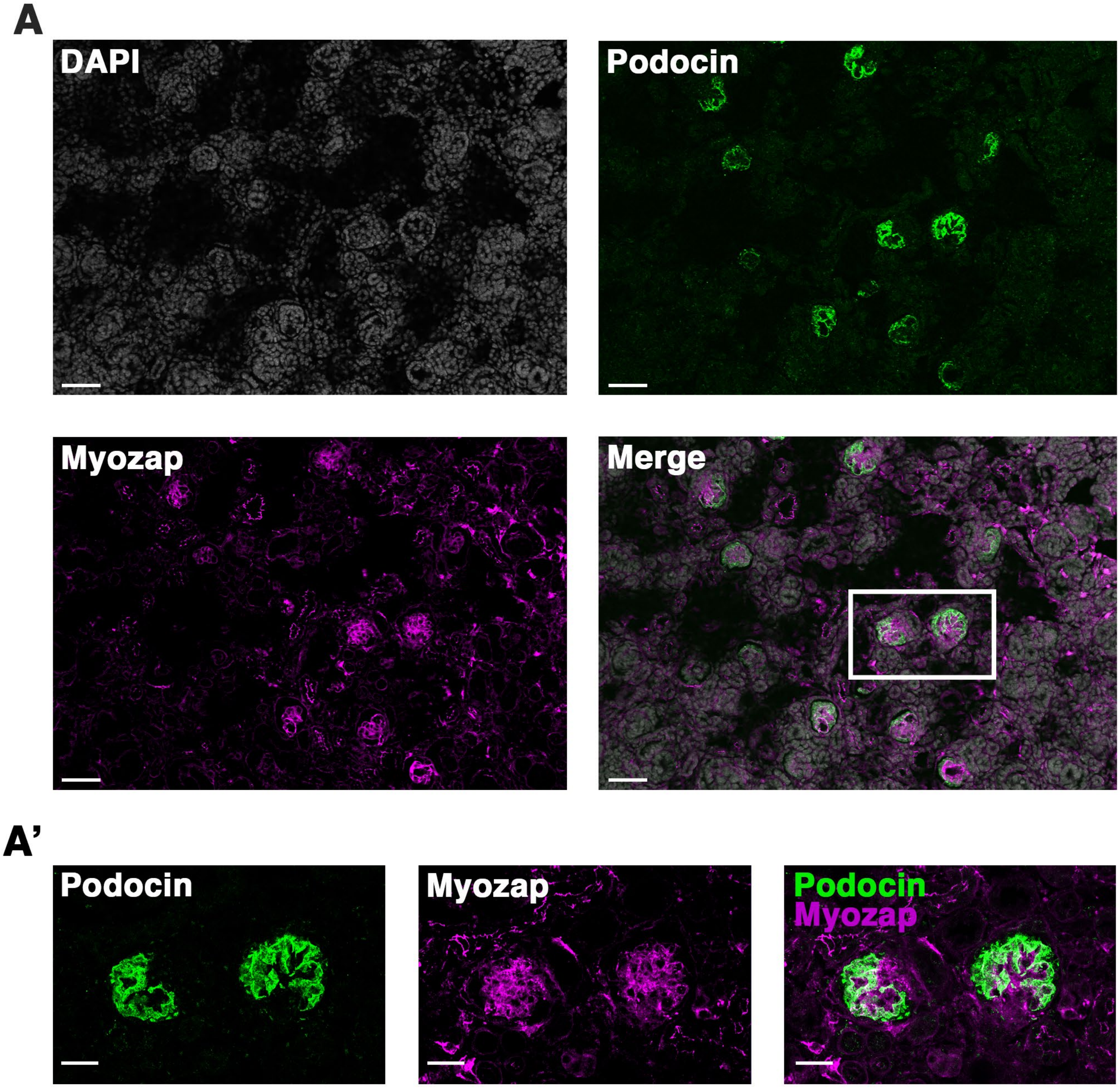


**Supplemental Figure 2. Myozap localizes to endothelium of glomeruli. (A)** Immunofluorescence analysis of Myozap (magenta) identifies strong localization to the central region of the glomerulus and endothelium outside the glomerulus. Proteomic profiling detected a significant Myozap signal (3.45 $Log_2$ FC), in two of three MS analyses. We observe some overlap (white) between Myozap (magenta) and podocin (green), while the majority of Myozap signal does not overlap with podocin and instead localizes to presumptive endothelial cells. Scale bar: 50 μm **(A')** Highlighted white boxed region from **(A)** indicating staining of Myozap within the endothelium of the glomerulus and some overlap with podocin. Scale bar: 20 μm.

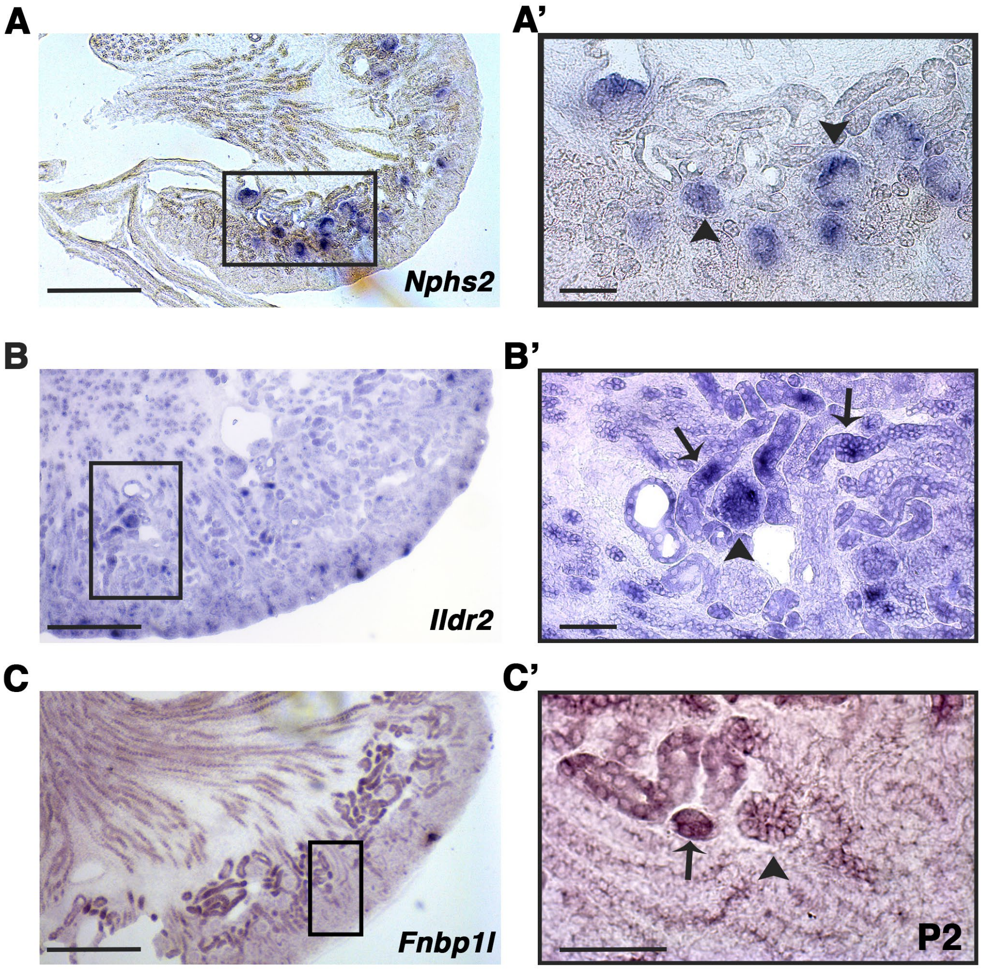


**Supplemental Figure 3. *In situ* hybridization confirms glomerular and some tubule cell expression of *Ildr2* and *Fnbp1l* in P2 kidneys.** P2 kidney sections hybridized with antisense riboprobes against *Nphs2* (podocin)**,** *Ildr2*, and *Fnbp1l*. **(A)** *Nphs2* (purple) displays strong expression specifically in glomeruli at P2. The rectangular box in **(A)** is enlarged in **(A')** to highlight expression of *Nphs2* only in glomeruli, arrowheads denote example glomeruli **(B)** *Ildr2* transcripts (purple) are identified within glomeruli and tubules. The rectangular box in **(B)** is enlarged in **(B')** to denote expression of *Ildr2* in glomeruli, arrowhead, and in some tubules, black arrow. **(C)** *Fnbp1l* (purple) is identified in both glomeruli and tubules. The block rectangular box in **(C)** is enlarged in **(C')** to denote expression of *Fnbp1l* in glomeruli, arrowhead, and tubules, arrow. Scale bars in A–C: 500 μm. Scale bars in A'-C': 100 μm.

A

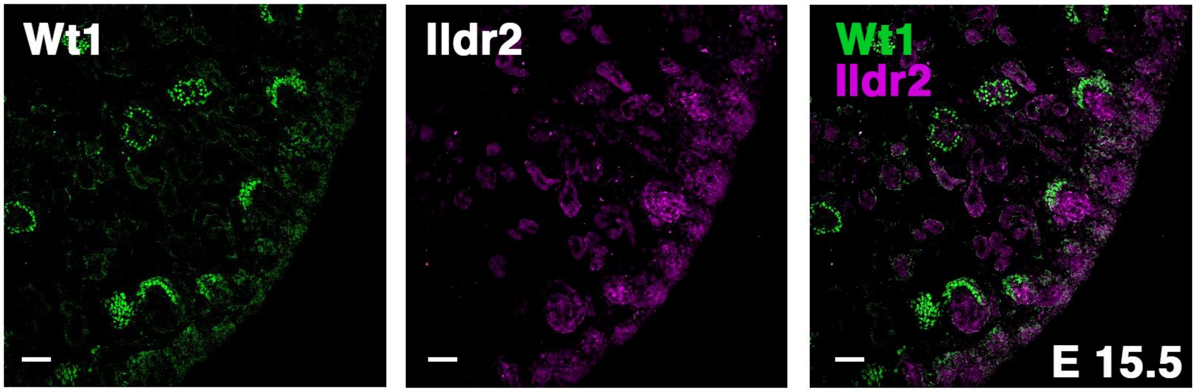


**Supplemental Figure 4. Ildr2 is detected in developing nephron structures.** Immunofluorescence analysis of E15.5 kidney sections co-immunostained for Ildr2 (magenta) Wt1 (podocytes, green). Scale bar: 50 μm.

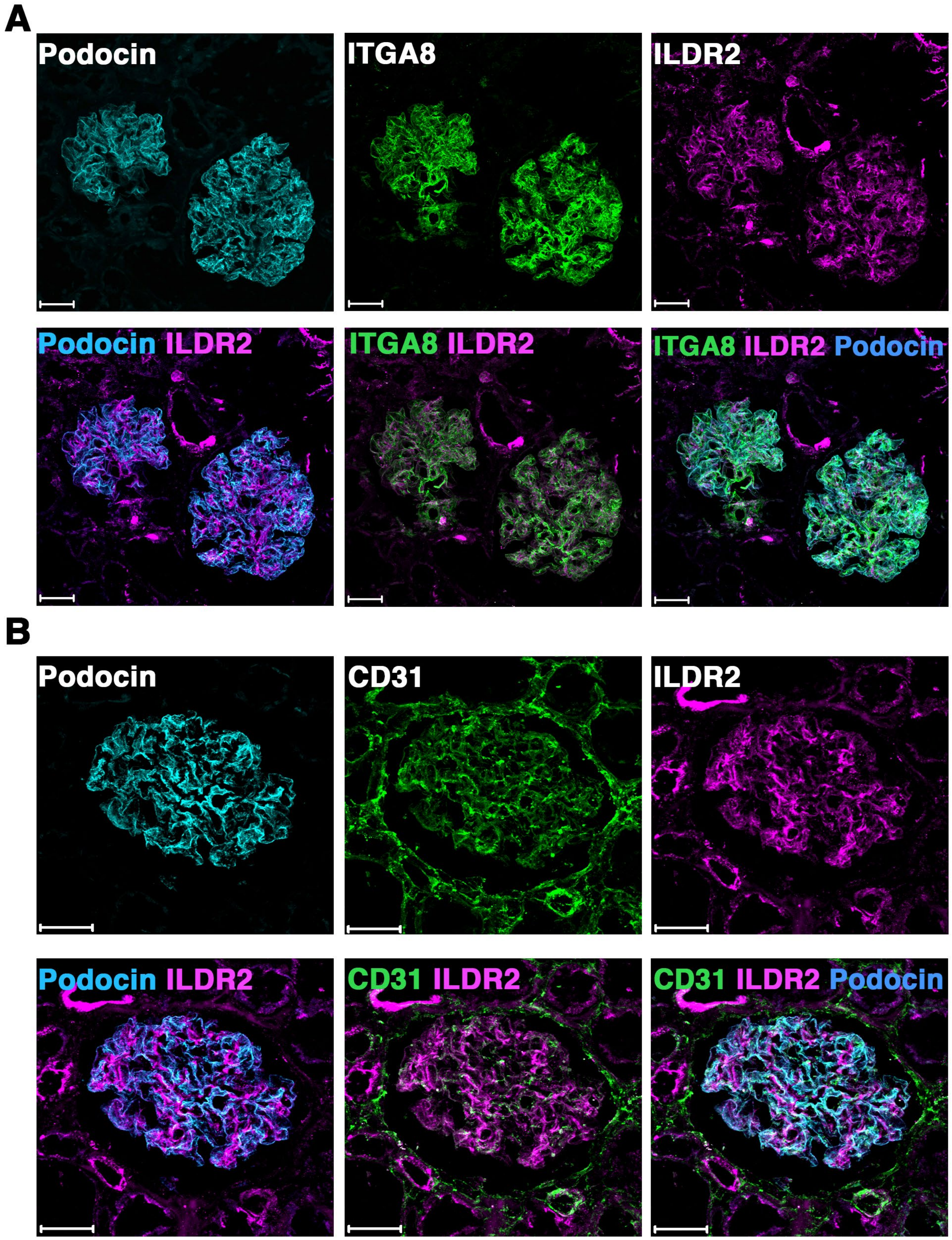


**Supplemental Figure 5. ILDR2 increases in detection, specifically in podocytes, of an aged human kidney.** Immunofluorescence analysis of sections from 91yo human kidney tissue identifies an increase in ILDR2 signal specifically within podocytes. **(A)** Colocalization of ITGA8 (mesangium, green), ILDR2 (magenta), and podocin (cyan) **(B)** Colocalization of CD31 (endothelium, green), ILDR2 (magenta), and podocin (cyan). Scale bar: 50 μm.
